# POGZ safeguards neuronal gene chromatin architecture and transcription

**DOI:** 10.64898/2026.07.02.736104

**Authors:** Natasha Ann Mariano, Katerina J. Williams, Katie Munechika, Katherine M. Bonefas, Yu Sun, Weiyu Zhang, Jared L. Klein, Kevin Monahan, Haiyuan Yu, Cedric Feschotte, Eirene Markenscoff-Papadimitriou

## Abstract

Disruption of chromatin organization is a common pathogenic mechanism in neurodevelopmental disorders, yet how changes in 3D genome architecture relate to transcriptional dysfunction in the developing brain remains unclear. POGZ, a transposase-derived chromatin regulator mutated in White–Sutton syndrome and autism spectrum disorder (ASD) has been linked to both heterochromatin and accessible regulatory DNA, but its in vivo function in the brain is unresolved. Using immunoprecipitation–mass spectrometry in embryonic day 13.5 (E13.5) mouse cortex, we identify the H3K9 methyltransferases G9a/GLP as principal POGZ interactors, placing POGZ within the core H3K9 methylation machinery in vivo. We find that POGZ loss in embryonic mouse cortex drives bidirectional, megabase-scale redistribution of H3K9me3, with ectopic losses and gains over discrete neuronal gene loci. In developing *Pogz^-/-^*cortex, regions with H3K9me3 gains are repositioned to the nuclear lamina and exhibit strengthened B-compartment scores by Micro-C, locus-restricted erosion of TAD architecture, weakened boundary insulation, and reduced CTCF occupancy. Within these domains, nascent RNA synthesis at neuronal genes is markedly diminished. These results identify POGZ as a G9a/GLP-associated chromatin regulator that protects neurodevelopmental gene domains from heterochromatinization and perinuclear sequestering, preserving 3D architecture and transcription during cortical development.

**Major Points:**

- POGZ interacts with the G9a/GLP H3K9 methyltransferase complex in the developing mouse cortex, identified by IP–mass spectrometry.
- Loss of POGZ drives megabase-scale redistribution of H3K9me3 and increases compartmentalization and nuclear-lamina association of neurodevelopmental gene loci, shown by ChIP-seq, Micro-C, and DNA FISH.
- Micro-C reveals disrupted TAD architecture at specific neuronal gene loci in Pogz⁻/⁻.PRO-seq shows POGZ is required to maintain nascent RNA synthesis of neurodevelopmental genes within disrupted chromatin domains.

## Introduction

*Pogo* transposable element derived with ZNF domain (POGZ) is among the most frequently mutated genes in individuals with autism spectrum disorder (ASD)(De Rubeis *et al*., 2014; Iossifov *et al*., 2014; Satterstrom *et al*., 2020), intellectual disability (ID)(Stessman *et al*., 2016), and developmental disorders (Deciphering Developmental Disorders Study, 2015), and is the defining gene of White–Sutton syndrome, a neurodevelopmental disorder (NDD) characterized by cognitive impairment, autism-like behaviors, and craniofacial abnormalities (Batzir *et al*., 2020). Large-scale sequencing studies consistently implicate POGZ as a high-confidence NDD risk gene, yet the molecular functions of POGZ in the mammalian brain remain incompletely understood.

POGZ is a highly conserved gene across vertebrates that evolved by combining exons encoding multiple zinc fingers (ZNF domain) with a complete transposase domain derived from an ancestral *Pogo*-like transposable element, resulting in a ∼1400 amino acid chimeric protein (Cosby *et al*., 2021). Following its link to NDDs, several studies have investigated the molecular functions of POGZ in human cell lines. Notably, the N-terminal ZNF domain of POGZ has been shown to mediate binding to heterochromatin protein 1 (HP1) isoforms (Nozawa *et al*., 2010). Subsequent work showed that POGZ forms separate nuclear complexes together with factors such as CHAMP1 (Li *et al*., 2022) and ADNP (Markenscoff-Papadimitriou *et al*., 2021), also linked to NDDs, and has been reported to promote heterochromatin formation at pericentromeric regions and transposable elements, as well as homology-directed repair (Heath *et al*., 2022; Sun *et al*., 2023; Li *et al*., 2025). These findings support a model in which POGZ participates in the establishment or maintenance of heterochromatic domains. Notably, many of these mechanistic insights derive from transformed cell lines or *in vitro* systems, leaving unresolved how POGZ functions within the chromatin landscape of the developing brain in vivo. POGZ is also linked to promoting accessibility at enhancers neighboring neurodevelopmental genes by recruiting esBAF/SWI–SNF chromatin remodeling complexes in neural progenitors (Sun, Cheng and Sun, 2022). Thus, POGZ has been connected both to the promotion of heterochromatin as well as to the maintenance of accessible, transcriptionally permissive euchromatin. These paradoxical activities suggest that POGZ functions as a context-dependent architectural protein involved in maintaining neuronal chromatin homeostasis.

Here, we set out to further characterize POGZ’s molecular function in the developing mouse cortex. We generated a Pogz-HA knock-in mouse to identify the POGZ protein interactome at embryonic day 13.5 (E13.5) by mass spectrometry, recovering previously reported interactors as well as new chromatin regulators including the H3K9 methyltransferases G9a and GLP. This finding places POGZ within the core H3K9 methylation machinery in vivo. We used a previously generated *Pogz* knockout mouse to assess the contribution of POGZ to the H3K9me3 heterochromatin landscape, 3D genome organization, and transcriptional activity of developing cortical neurons and integrated these data with new POGZ CUT&RUN binding maps in E13.5 cortex. We find that POGZ loss leads to the redistribution of H3K9me3 across mega-base scale regions together spanning ∼10% of the mouse genome: H3K9me3 is lost at 47 gene clusters including odorant receptors and clustered protocadherin genes, while H3K9me3 is gained at 49 domains harboring genes encoding key synaptic signaling molecules. Domains with H3K9me3 gains are repositioned to the nuclear lamina in *Pogz*⁻^/^⁻ cortex, exhibit reorganized genome compartments and weakened TADs, and show reduced nascent RNA synthesis at affected gene bodies. Together, our results link POGZ loss to coordinated, domain-scale changes in 3D genome organization, heterochromatin deposition, and transcriptional activity at a defined subset of neuronal loci, identifying POGZ as a chromatin regulator that safeguards neurodevelopmental gene expression during cortical development.

## Results

### POGZ interacts with the G9a/GLP H3K9 methyltransferase complex in the developing mouse cortex

To identify POGZ-interacting proteins in the developing brain, we generated a Pogz-HA knock-in mouse in which a C-terminal hemagglutinin (HA) epitope tag was inserted at the endogenous Pogz locus by CRISPR-Cas9 (Fig. S1A)(see Methods). HA-tagged POGZ was expressed at expected levels by western blot and is absent in wild-type controls (Fig. S1A). We performed anti-HA immunoprecipitation followed by mass spectrometry (IP-MS) on individually dissected E13.5 cortices from Pogz-HA and wild-type littermates (n=4, each genotype), see Methods.

Gene ontology analyses of proteins pulled down in our assay (log2fc >0.5) showed enrichment for terms such as heterochromatin formation (7 proteins) and DNA repair (7 proteins)(Fig. S1B). Among the most significantly enriched interactors were the H3K9 methyltransferases EHMT2 (G9a) and EHMT1 (GLP)( adj p-value <0.05 and log2FC>1) which form an obligate heterodimer responsible for the bulk of H3K9 mono-and dimethylation (Tachibana *et al*., 2005)(Fig. 1A). We also recovered the previously reported POGZ partner CHAMP1 (Li *et al*., 2025), additional chromatin-associated factors (HDGFL2, PRDM10, ZMYM4, ZNF428), and the Ku70/Ku80 DNA repair heterodimer (XRCC5/XRCC6). Consistent with prior work, we recovered Cbx3/HP1γ (adj. p = 0.0019), a heterochromatin scaffolding protein shown to interact with POGZ and co-occupy thousands of genomic loci in developing human and mouse cortex (Nozawa *et al*., 2010; Markenscoff-Papadimitriou *et al*., 2021).

**Figure 1.**
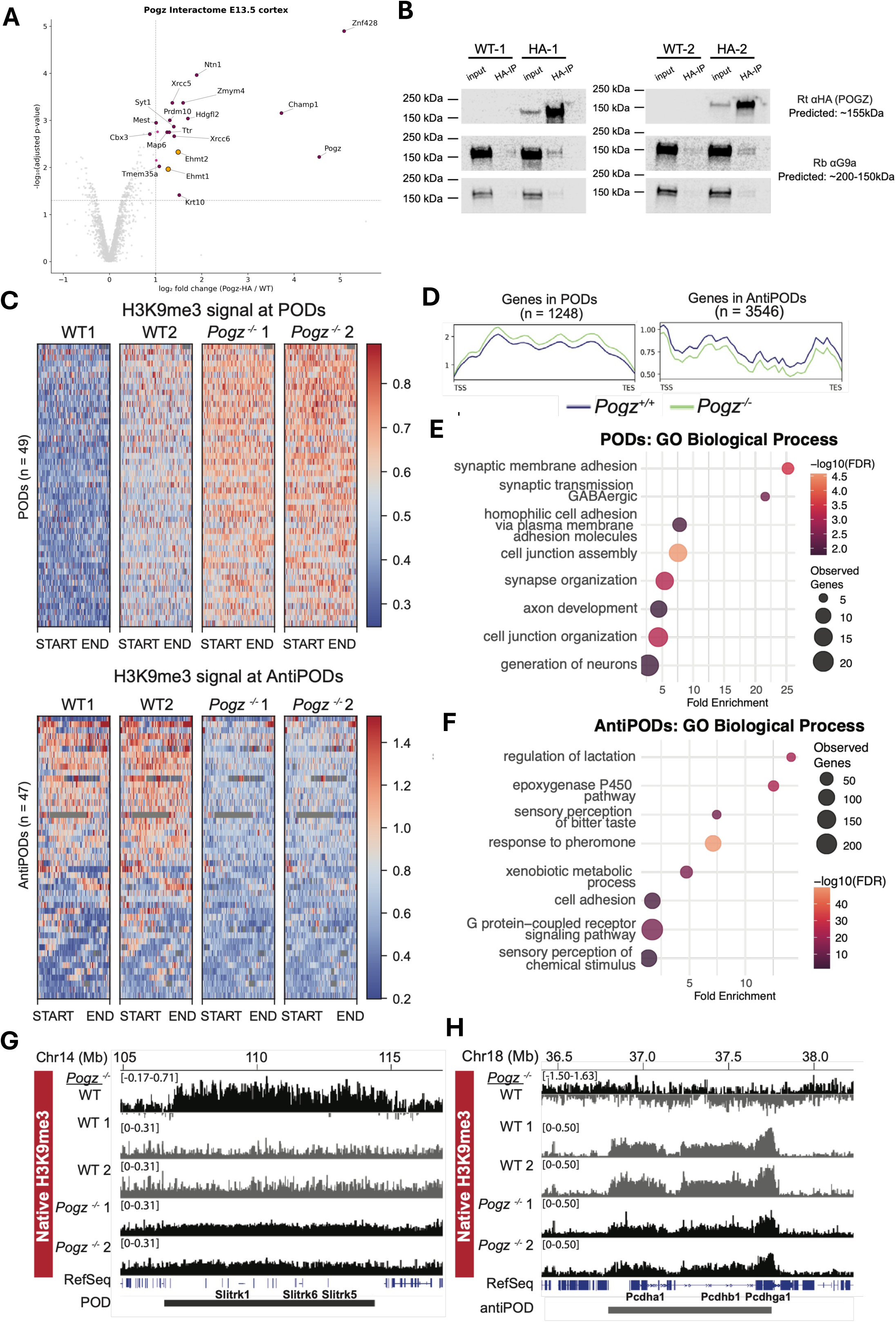
POGZ interacts with the G9a/GLP H3K9 methyltransferase complex and protects neuronal gene domains from heterochromatinization. (A) Volcano plot of proteins co-immunoprecipitated with anti-HA from E13.5 *Pogz-HA* knock-in cortex relative to wild-type controls (IP–MS, n = 4 biological replicates per genotype). x-axis, log₂ fold change (*Pogz-HA* / WT); y-axis, –log₁₀ adjusted *p*-value. Dashed lines mark significance thresholds (adj. *p* < 0.05, log₂FC > 1). The H3K9 methyltransferases EHMT2 (G9a) and EHMT1 (GLP) are highlighted in orange; other significantly enriched interactors are labeled, including CHAMP1, Cbx3/HP1γ, and the Ku70/Ku80 (XRCC6/XRCC5) heterodimer. (B) Co-immunoprecipitation of POGZ and G9a in the developing mouse cortex. HA and G9a expression in wild-type (WT-1, WT-2) and *Pogz*-HA heterozygous (HA-1, HA-2) cortices, with 5% input (input, 20 ug total protein) or after HA pull down (HA-IP). G9a blots shown are the same blots with higher and lower exposure settings. (C) Heatmaps of native H3K9me3 ChIP-seq signal in two wild-type (WT1, WT2) and two *Pogz⁻ᐟ⁻* (1, 2) E13.5 cortices, plotted across length-scaled domains (START–END) for PODs (top, n = 49) and anti-PODs (bottom, n = 47). PODs gain H3K9me3 in *Pogz⁻ᐟ⁻*; anti-PODs lose H3K9me3. (D) Aggregate H3K9me3 ChIP-seq signal in wild-type and *Pogz⁻ᐟ⁻* cortex over RefSeq genes within PODs (n = 1,248) and anti-PODs (n = 3,546). (E, F) GO enrichment of genes located within PODs (E) and anti-PODs (F). Dot size, gene count; color, significance; x-axis, fold enrichment. (G) Native H3K9me3 ChIP-seq tracks across the *Slitrk1/6/5* POD (chr14, ∼105–116 Mb). Top, *Pogz⁻ᐟ⁻* normalized to WT track (black); below, WT, WT1, WT2, *Pogz⁻ᐟ⁻* 1, and *Pogz⁻ᐟ⁻* 2. replicates. RefSeq genes and POD interval are shown below. H3K9me3 is broadly increased across the domain in *Pogz⁻ᐟ⁻*. (H) As in (G) for the clustered protocadherin (*Pcdhα/β/γ*) anti-POD (chr18, ∼36.5–38 Mb), showing reduced H3K9me3 across the domain in *Pogz⁻ᐟ⁻*.

Whereas earlier studies placed POGZ within the heterochromatin scaffolding machinery through its association with HP1 isoforms, our in vivo interactome analysis reveals that POGZ also engages the H3K9 methyltransferase writers G9a and GLP directly. G9a has been shown to be crucial for positive regulation of olfactory receptor genes in olfactory sensory neurons (Lyons *et al*., 2014) and for silencing of non-neuronal genes in cortical neurons (Schaefer, 2012). This positions POGZ at the core of the H3K9 methylation machinery in the developing cortex. We confirmed the POGZ–G9a interaction by co-immunoprecipitation in E13.5 cortex (Fig. 1B). This interaction motivated us to ask how POGZ shapes the H3K9 methylation landscape in vivo.

### Loss of POGZ induces bidirectional, megabase-scale redistribution of H3K9me3 in the developing cortex

G9a/GLP catalyze mono and dimethylation of H3K9, which serve as the preferred substrates for SUV39H-mediated H3K9me3, the dominant heterochromatic state in differentiated cells (Padeken, Methot and Gasser, 2022). We therefore profiled H3K9me3 in a previously characterized Pogz constitutive knockout mouse (Markenscoff-Papadimitriou *et al*., 2021). *Pogz^-/-^* embryos are perinatally lethal but undergo grossly normal early corticogenesis: prior analyses at E13.5 show no significant alterations in cortical progenitor or neuronal populations, cortical plate formation, or layer-specific marker expression, arguing against major changes in cell-type composition at this stage. Heterozygous *Pogz* mice are viable and exhibit autism-related behavioral phenotypes (Cunniff *et al*., 2020). We performed native ChIP-seq with an anti-H3K9me3 antibody on nuclei isolated from individual E13.5 cortices from wildtype and *Pogz*^-/-^ littermates, an approach used for assaying the H3K9me3 landscape in neurons without crosslinking or sonication. Libraries met all quality control metrics, and principal component analysis confirmed strong clustering by genotype with high reproducibility between biological replicates (Fig. S2B-C). To identify regions of differential H3K9me3 deposition, we performed differential domain analysis using SICER, which detects broad heterochromatin changes at the megabase-scale (Xu *et al*., 2014) (see Methods).

Loss of POGZ resulted in the megabase-scale redistribution of H3K9me3 across the embryonic cortex. Using SICER, we identified 49 large genomic domains (mean length 3.3 megabases (Mb)) with increased H3K9me3 signal, and 47 genomic domains (mean length 2.2 Mb) with reduced H3K9me3 signal in *Pogz^-/-^* cortex relative to wildtype (Fig. 1C). We refer to regions that gained H3K9me3 as Pogz-dependent domains (PODs), which together span 6% of the mm10 genome, and regions that lost H3K9me3 as anti-PODs, spanning 3.9% of the genome. The “anti-” designation denotes the opposite direction of H3K9me3 change, not a lesser dependence on POGZ, as both domain classes arise directly from POGZ loss of function. Collectively, Pogz-dependent changes in H3K9me3 levels affect ∼10% of the genome in the mouse cortex.

Genes in PODs (n=1,248) exhibit gains in H3K9me3 levels in *Pogz^-/-^*, while genes in anti-PODS (n=3,546) show decreases in H3K9me3 (Fig. 1D). POD genes were significantly enriched for neurodevelopmental gene ontology (GO) terms, including “synaptic membrane adhesion,” “homophilic cell adhesion,” “axon development,” and “generation of neurons” (Fig.1E). In contrast, anti-PODs were associated with a broader and more heterogeneous set of GO terms. These included “response to pheromone,” reflecting vomeronasal receptor gene clusters; “G protein–coupled receptor signaling pathway,” driven primarily by olfactory receptor gene loci; and “sensory perception of chemical stimulus” (Fig. 1F).

An example POD locus is on chromosome 14 at the *Slitrk1,6,5* gene cluster (Fig. 1G). Slit and Trk-like family member (*Slitrk*) genes encode transmembrane proteins essential for neuronal development, specifically in guiding neurite outgrowth and synapse formation in the brain, and *Slitrk1* has been linked to Tourette’s disorder (Abelson *et al*., 2005; Song *et al*., 2015). At this locus we observe increased levels of H3K9me3 across the 7.8 Mb-long POD encompassing the *Slitrk* cluster. An example anti-POD locus is the 1-Mb locus containing the clustered protocadherin genes (*Pcdhgα,β,γ*) (Fig. 1H). H3K9me3 heterochromatin at this locus has been previously shown in olfactory sensory neurons to be compatible with variable protocadherin isoform selection (Kiefer *et al*., 2023). Further example POD and anti-POD loci at neuronal genes are shown in Figures S2D and S2E.

In sum, POGZ deletion induces megabase-scale remodeling of H3K9me3, increasing heterochromatinization at a set of neuronal gene–containing regions while simultaneously reducing H3K9me3 at other neuronal gene containing regions of the genome.

### H3K9me3 redistribution in Pogz knockout is coupled to localized changes in 3D compartments

The megabase-scale gains and losses of H3K9me3 observed following *Pogz* knockdown suggested that Pogz may influence chromatin organization beyond local regulatory elements, potentially affecting higher-order genome folding. To assess whether *Pogz* loss alters higher-order organization, we performed Micro-C on E13.5 cortical tissue from *Pogz*⁺^/^⁺ and *Pogz*⁻^/^⁻ embryos and libraries were generated from individual cortices (see Methods). Replicates clustered by genotype in principal component analysis (PCA) of A/B compartment scores, while cis/trans interactions showed that they are similar for all libraries, and correlation scores between replicates had a R^2^ value >0.98, indicating high data quality and reproducibility (Figs S3A-D). “A” compartments are generally associated with euchromatin, whereas “B” compartments are associated with heterochromatin (Lieberman-Aiden *et al*., 2009). Genome-wide compartment identity was largely preserved in *Pogz^-/-^*, with no significant changes associated with A-to-B or B-to-A compartment switches (Fig. 2A).

**Figure 2.**
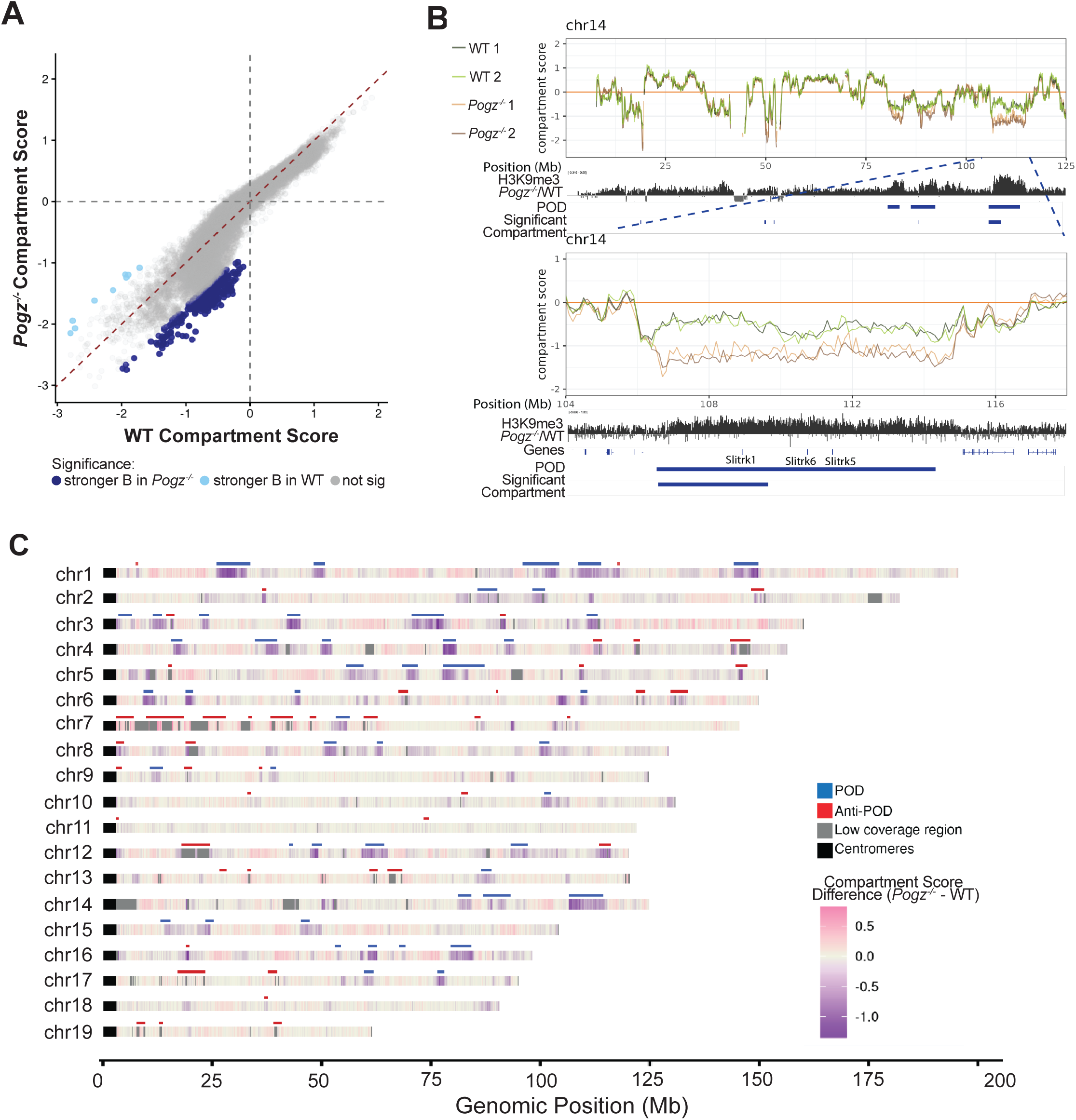
Compartment score changes in *Pogz⁻ᐟ⁻* cortex. **(A)** Micro-C compartment strength scores in wild-type (x-axis) versus *Pogz⁻ᐟ⁻* (y-axis) E13.5 cortex (100-kb bins, merged replicates). Pearson r = 0.944. Points colored by paired Z-test (adj. *p* < 0.1): non-significant (gray), significant B score strength change in *Pogz⁻ᐟ⁻* (dark blue), significant B score strength change in wild-type (light blue) **(B)** Chromosome 14 tracks: per-replicate compartment strength scores for *Pogz⁻ᐟ⁻* (ko1, ko2) and wild-type (wt1, wt2); H3K9me3 *Pogz⁻ᐟ⁻*/WT signal; POD intervals; significantly changed A/B regions; RefSeq genes. Top, full chromosome; bottom, zoom into the *Slitrk1/6/5* POD. **(C)** Genome-wide compartment-score change (*Pogz⁻ᐟ⁻* − WT) across autosomes (chr1–chr19) by genomic position (Mb). Color, change in compartment score (CS). PODs (blue), anti-PODs (red), low-coverage regions (gray), and centromeres (black) annotated.

While chromatin compartment identity was stable, we observed many discrete, megabase-scale loci with significant shifts in B compartment score strength with POGZ loss (Fig. 2A). Paired Z test identified 469 loci showing significant (padj<0.1) changes in B compartment strength: 460 of these regions showed a significant change towards a stronger B compartment in the knockout, while 9 showed a significant change towards A compartment in the knockout (Fig. 2A, Fig. S3E). Thus, *Pogz* loss does not globally reorganize compartment identity but instead induces changes in compartment strength for a subset of large heterochromatic domains or B compartments.

Mapping compartment score changes across the genome revealed a strong correspondence with POGZ-dependent H3K9me3 domains (PODs)(Fig. 2B-C, Fig. S4). In contrast, anti-PODs appear to show little detectable compartmental changes (Fig. 2C, Fig. S4), but this is difficult to assess due to low sequencing coverage in the Micro-C data likely caused by very low chromatin accessibility in these regions. Nonetheless, along chromosomes we generally observe that regions exhibiting gains in H3K9me3 corresponded to strengthened B-compartment scores, whereas anti-PODs correspond with decreased B-compartment strength (Fig. 2B, Fig. S4).

Consistent with these observations, both Fisher’s exact and permutation test revealed a highly significant association between PODs and regions exhibiting changes in B-compartment strength (Fisher’s pval = 5.14e-254, permutation p = 1.00e-3). Together, these results indicate that POGZ knockout leads to coordinated alterations in H3K9me3 deposition and large-scale chromatin compartmentalization.

### Neurodevelopmental gene loci in PODs are repositioned to the nuclear lamina in *Pogz⁻^/^⁻*cortex

The strengthened B-compartment scores at PODs predicted that these loci should also reorganize within the nucleus, since B-compartment chromatin preferentially associates with the nuclear lamina (Lieberman-Aiden *et al*., 2009; van Steensel and Belmont, 2017). To test this directly, we performed DNA fluorescent *in situ* hybridization (FISH) combined with Lamin B1 immunofluorescence on coronal cryosections of E13.5 wild-type and *Pogz^-/-^* cortex (Fig. 3A-C). We selected BAC probes spanning three POD gene clusters — *Adgrl3*, *Flrt2*, and *Slitrk5* — as well as the β-actin locus as control, and scored each allele as lamina-associated if its FISH signal directly overlapped the Lamin B1 ring (Fig. 3D-G, 3D’-G’, S5A-D). Scoring included all DAPI-positive nuclei spanning the ventricular zone to the cortical plate, encompassing both progenitors and post-mitotic neurons in E13.5 cortex, respectively (see supplementary methods).

**Figure 3.**
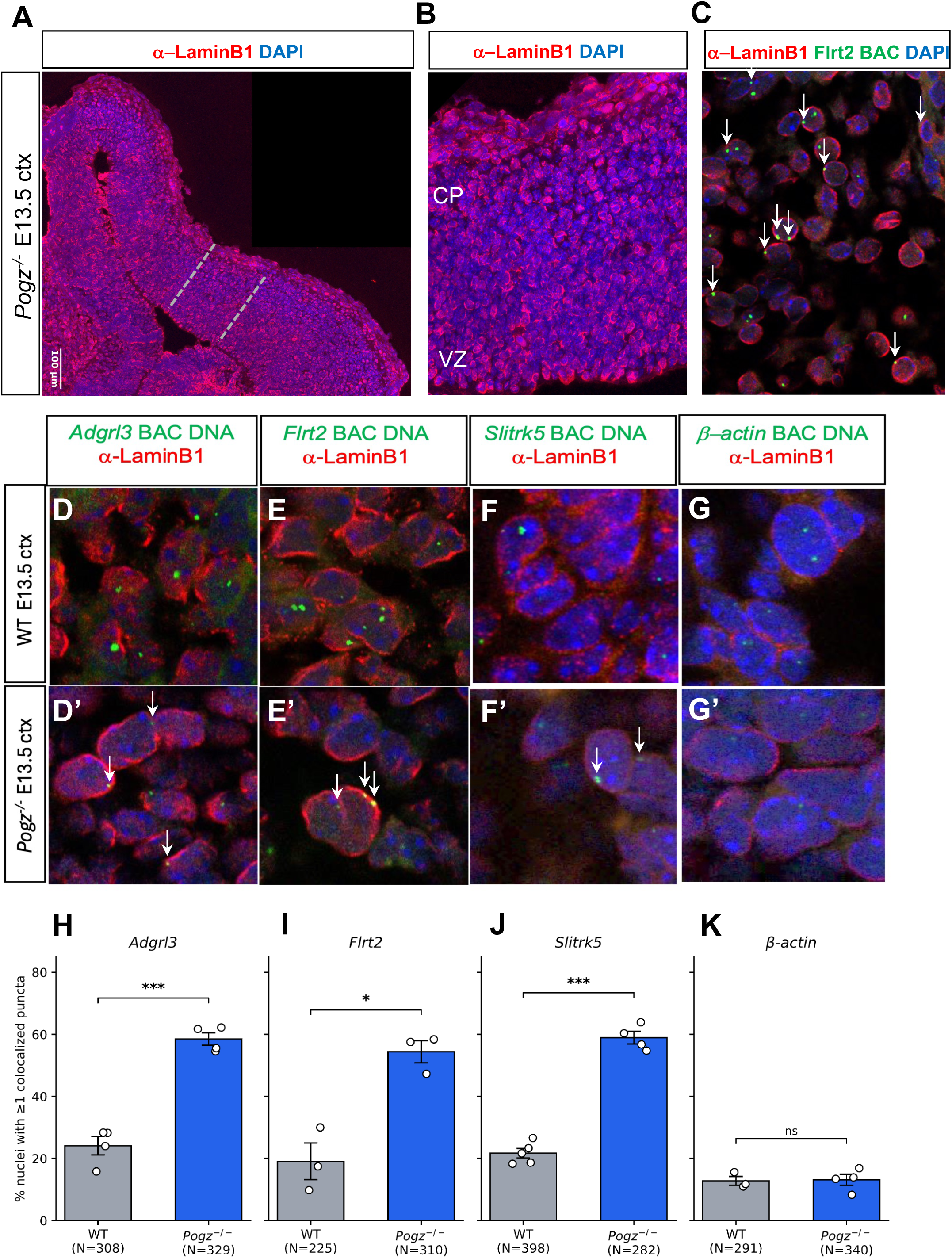
POD loci reposition to the nuclear lamina in *Pogz⁻ᐟ⁻* cortex. **(A–C)** Representative E13.5 *Pogz⁻ᐟ⁻* cortical sections stained for Lamin B1 (red) and DAPI (blue), shown alone (A), at higher magnification at dotted inset (B), and combined with *Flrt2* BAC DNA FISH (green) (C), single z-plane shown. Cortical plate (CP) and ventricular zone (VZ) labeled where cortical projection neurons and progenitors, respectively, reside.Arrows indicate *Flrt2* DNA FISH puncta co-localized with Lamin-B1. **(D–G)** DNA FISH (green) for POD loci *Adgrl3* (D), *Flrt2* (E), and *Slitrk5* (F), and the β-actin control locus (G), combined with Lamin B1 immunofluorescence (red) and DAPI (blue) in wild-type E13.5 cortex. **(D′–G′)** Same probes in *Pogz⁻ᐟ⁻* cortex. Arrows indicate DNA FISH puncta co-localized with Lamin-B1; single z-plane shown. **(H–K)** Quantification of the percentage of nuclei with ≥1 FISH punctum colocalized with the Lamin B1 signal in wild-type (gray) and *Pogz⁻ᐟ⁻* (blue) cortex, for *Adgrl3* (H), *Flrt2* (I), *Slitrk5* (J), and β-actin (K). Bars, mean ± SEM; points, individual embryos; N nuclei per genotype indicated below. Two-tailed Welch’s *t*-test: *, *p* < 0.05; ***, *p* < 0.001; ns, not significant.

In wild-type cortex, POD alleles were rarely positioned at the nuclear periphery (∼20% of nuclei with ≥1 lamina-contacting allele, N = 931 nuclei from 12 wildtype embryos)(Fig. 3H-J), consistent with these loci residing in nuclear-interior chromatin and being excluded from canonical lamina-associated domains (LADs). In *Pogz^-/-^* cortex, all three POD loci were repositioned to the nuclear lamina, with ∼60% of nuclei harboring a lamina-contacting allele at each of the three loci (p <0.05, two-tailed Welch’s t-test; N= 921 nuclei from 11 *Pogz^-/-^*embryos)(Fig. 3H-J); β-actin co-localization with the nuclear lamina was unchanged (Fig. 3K). Together, these single-cell imaging data are congruent with the changes in compartment scores inferred from Micro-C data and the changes in H3K9me3, and converge to the notion that POGZ is required to keep neurodevelopmental gene clusters away from the repressive nuclear periphery during cortical development.

### POGZ binds regulatory elements at neurodevelopmental gene loci in the developing cortex

To map POGZ binding genome-wide in vivo and connect it to the architectural and heterochromatin changes observed in Pogz^⁻/⁻^ cortex, we performed CUT&RUN (Skene and Henikoff, 2017) on individual E13.5 cortices from wild-type embryos, using *Pogz^-/-^* littermates and IgG as controls. Libraries met all quality control metrics (Fig. S6A,B), and signal was specific: POGZ binding profiles showed strong enrichment in wild-type cortex with negligible signal in *Pogz^-/-^* and IgG controls (Fig. S6C,D), confirming antibody specificity in vivo. At the Follistatin-like 5 (*Fstl5)* locus, multiple POGZ peaks were detected in wild-type cortex and were absent in *Pogz^-/-^* and IgG samples (Fig. 4A), illustrating the high signal-to-noise of the assay at individual loci.

**Figure 4.**
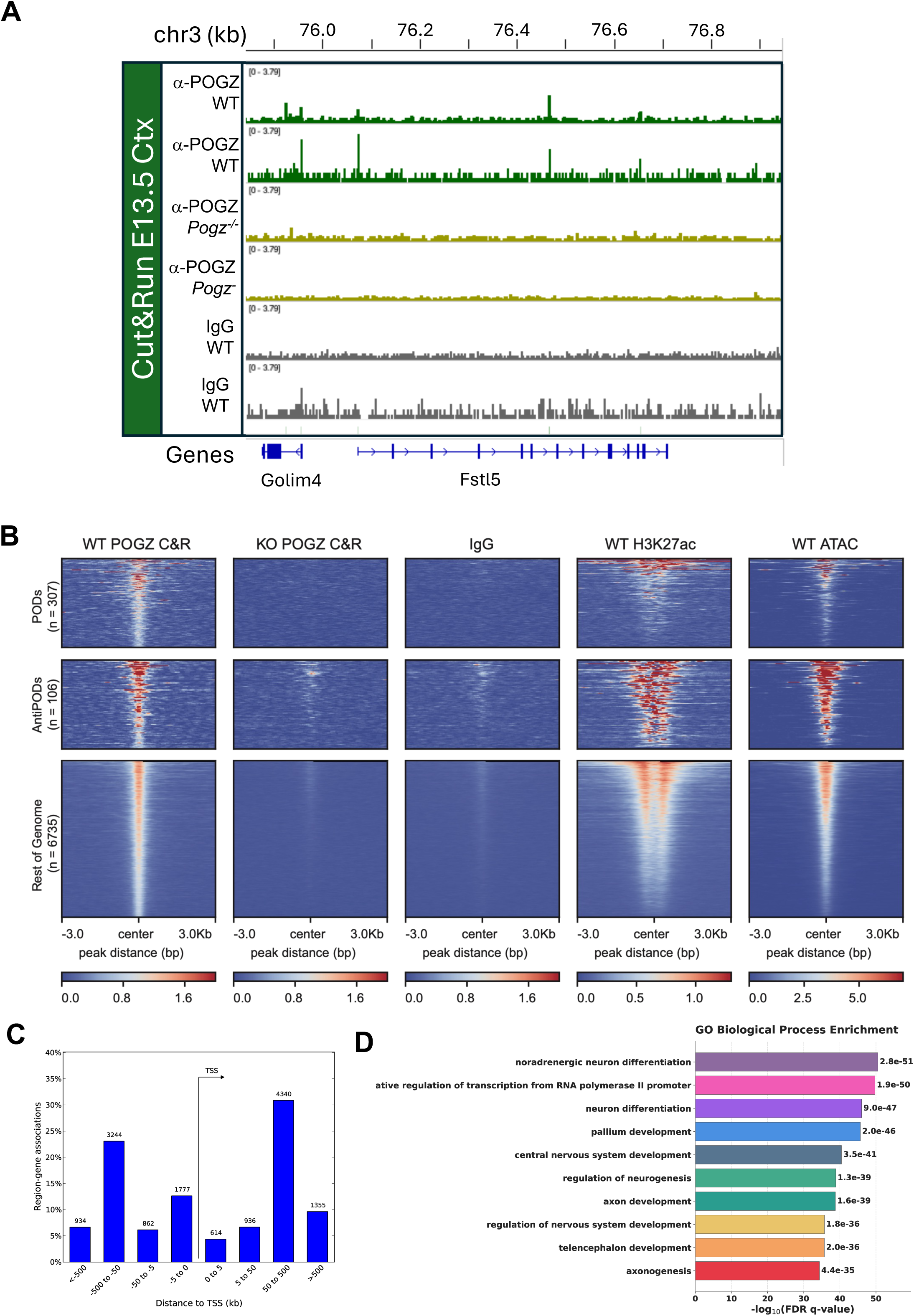
POGZ binds open chromatin at active regulatory elements at PODs. **(A)** Genome browser tracks of POGZ CUT&RUN signal in merge wild-type (α-POGZ WT) and *Pogz⁻ᐟ⁻* (α-POGZ *Pogz⁻ᐟ⁻*) E13.5 cortex, with IgG control, at the *Golm4/Fstl5* POD locus. RefSeq genes below. **(B)** Heatmaps of WT POGZ CUT&RUN, *Pogz⁻ᐟ⁻* POGZ CUT&RUN, IgG, WT H3K27ac, and WT ATAC-seq signal, centered on POGZ peaks (±3 kb), at PODs (n = 307), anti-PODs (n = 106), and the rest of the genome (n = 6,735). **(C)** Distribution of POGZ CUT&RUN peaks relative to annotated transcription start sites (TSS). **(D)** GO Biological Process enrichment of genes proximal to POGZ peaks (GREAT). Bar length, fold enrichment; *p*-values indicated.

Using MACS3 with *Pogz^-/-^* as background, we identified 7,148 high-confidence POGZ binding sites in E13.5 cortex (see Methods). POGZ CUT&RUN signal was observed at these peaks across PODs, anti-PODs, and the broader genome, with negligible signal in knockout controls at all three categories (Fig. 4B, S6C). POGZ peaks were strongly enriched for marks of active regulatory chromatin: signal coincided with open chromatin defined by ATAC-seq and with H3K27ac, a histone modification of active enhancer and promoter elements (Markenscoff-Papadimitriou *et al*., 2021)(Fig. 4B). Consistent with this, genomic distribution analysis using GREAT showed that POGZ binding is predominantly distal and intergenic, but a substantial fraction of peaks (32.5%) map within 5kb of annotated transcription start sites (Fig. 4C)(McLean *et al*., 2010). Thus, POGZ binds preferentially within “open” chromatin regions (OCRs) in the genome, including but not restricted to a subset of promoter regions, and with a subset of binding sites lying within the heterochromatic PODs and anti-PODs. Genes proximal to POGZ binding sites were significantly enriched for neurodevelopmental gene ontology terms, including “axon development,” “synapse organization,” “regulation of neurogenesis,” and “pallium development” (Fig. 4D). Together, these data establish POGZ as a regulator of accessible, distal and proximal regulatory elements at neurodevelopmental gene loci in the developing cortex.

To assess whether POGZ binding is enriched at PODs, we compared the overlap between POGZ-bound OCRs within and outside of PODs. In wild-type cortex, we found that 31% of OCRs within PODs were POGZ-bound, compared to 13.5% of OCRs genome-wide, a 2.3-fold significant enrichment (odds ratio = 2.88, 95% CI 2.49–3.33, p < 2.2 × 10⁻¹⁶, Fisher’s exact test; Fig. S6E). In contrast, OCRs within anti-PODs were slightly depleted for POGZ binding relative to the genome-wide background (10.9% vs. 13.8%, odds ratio = 0.76, p < 0.01; Fig. S6F). Thus, POGZ preferentially occupies accessible regulatory DNA within the very domains that gain ectopic H3K9me3 when POGZ is lost, thereby linking POGZ binding to the loci most vulnerable to heterochromatinization.

To ask whether POGZ engages the same partners across all of its binding sites in vivo, we compared our POGZ CUT&RUN to ADNP occupancy in E13.5 cortex (Markenscoff-Papadimitriou et al., 2021), since POGZ and ADNP form the ChAHP complex (Ostapcuk *et al*., 2018). Across the genome, ADNP signal coincided strongly with POGZ peaks (Fig. S6G), consistent with POGZ acting within ChAHP at the majority of its binding sites during cortical development. Strikingly, however, this co-occupancy broke down within PODs and anti-PODs: at POGZ peaks inside these domains, ADNP signal was markedly reduced or absent, despite robust POGZ binding (Fig. S6G). Thus, POGZ engages ADNP/ChAHP at most of its genome-wide sites, but the POD and anti-POD loci where it specifically regulates H3K9me3 deposition appear to be bound by POGZ in a ChAHP-independent manner, pointing to a distinct mode of POGZ action at the domains it protects from or commits to heterochromatinization.

### TADs are weakened within PODs in *Pogz^-/-^* cortex

The dramatic reorganization of nuclear architecture and POD localization in the *Pogz* knockout suggested that POGZ might affect finer levels of the chromatin organization hierarchy. To determine whether the megabase-scale changes in H3K9me3 deposition observed in *Pogz*⁻^/^⁻ cortex are accompanied by changes in higher-order chromatin organization, we examined chromatin folding at the level of topologically associating domains (TADs). TADs are self-interacting genomic units that constrain enhancer–promoter communication and typically span kilobase-to megabase-scale regions.

In wild-type cortex, PODs generally span multiple TADs, as expected from their megabase-scale size, an organization that may compartmentalize the enhancers controlling the neurodevelopmental genes they contain. We first focused on the 7.8 Mb POD on chromosome 14 encompassing the *Slitrk1,5,6* gene cluster. In wild-type mouse embryonic cortex, Micro-C analysis shows this POD is subdivided into multiple discrete TADs with strong boundary insulation (Fig. 5A). In contrast, the *Pogz*⁻^/^⁻ cortex revealed a pronounced collapse of local TAD architecture, characterized by the loss of four adjacent domains and a marked reduction in intra-domain interactions at this locus, while neighboring TADs outside this locus remained intact. Boundaries associated with weakened TADs were lost in the knockout as well (Fig. 5A). A similar pattern was observed at the Leucine-rich repeat and fibronectin type III domain containing 5 (*Lrfn5*) containing POD, with collapse of two adjacent TADs and loss of the intra-TAD boundary (Fig. 5B).

**Figure 5.**
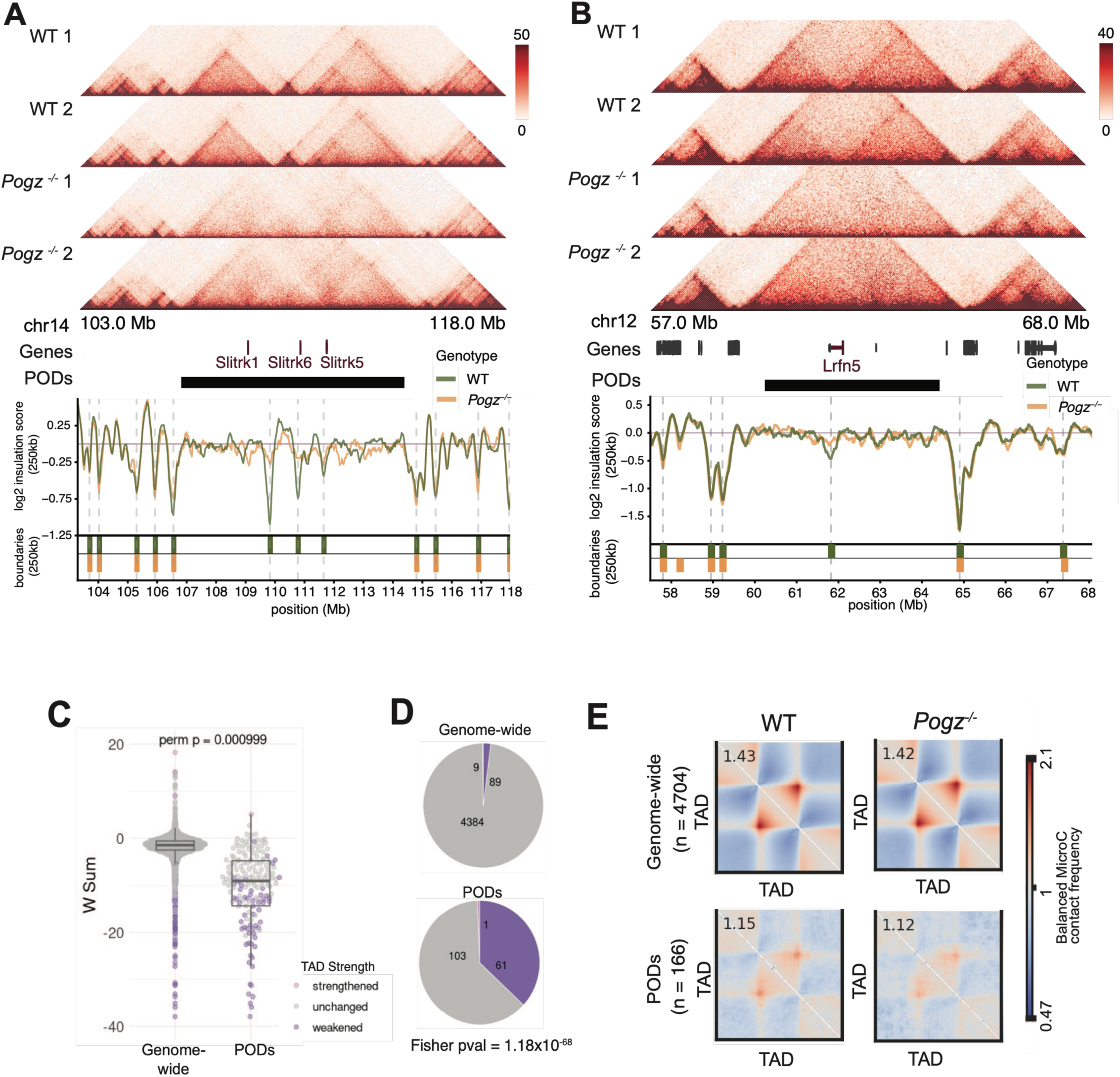
Localized disruptions in topologically associating domain (TAD) architecture in *Pogz⁻ᐟ⁻*. (A) Micro-C contact maps at the *Slitrk1/6/5* POD (chr14, 103.0–118.0 Mb) for two wild-type (WT1, WT2) and two *Pogz⁻ᐟ⁻* (1, 2) E13.5 cortices. RefSeq genes (*Slitrk1, Slitrk6, Slitrk5*) and POD interval shown below. Bottom: log₂ insulation score (250-kb window) for *Pogz⁻ᐟ⁻* (orange) and wild-type (green), with TAD boundary calls. (B) As in (A) for the *Lrfn5* POD (chr12). (C) Per-TAD interaction strength change (W sum) for TADs genome-wide versus within PODs. Points colored by classification: strengthened, unchanged, weakened. Permutation *p* = 1×10^3^. (D) Proportion of TADs classified as strengthened, unchanged, or weakened genome-wide (top) and within PODs (bottom). Fisher’s exact test *p* = 1.18 × 10⁻⁶⁸. (E) Aggregate Micro-C contact frequency over scaled TADs (±flanks) in wild-type and *Pogz⁻ᐟ⁻* cortex, genome-wide (n = 4,704) and within PODs (n = 166). Values, mean intra-TAD enrichment.

To determine whether these architectural changes at PODs reflect a global alteration in TAD organization or are restricted to specific genomic regions, we next performed genome-wide TAD calling using HiCExplorer on merged Micro-C datasets from each genotype (see Methods). We quantified changes in intra-TAD interaction strength between wildtype and *Pogz^-/-^* cortex. The collapsed TADs at the *Slitrk1,6,5* and *Lrfn5* loci were identified as significantly weakened in *Pogz^-/-^* cortex (Fig. 5A,B). Genome-wide, however, TAD organization was largely preserved: of 4,482 TADs identified, 4,384 remained unchanged, while only 98 were significantly weakened and 9 strengthened in *Pogz*⁻^/^⁻ cortex (p<0.05) (Fig. 5C,D). These results indicate that Pogz knockout does not broadly disrupt chromatin domain architecture.

Strikingly, 61 of the 98 TADs exhibiting weakened interaction strengths genome-wide were located within PODs, a highly significant enrichment (p=1.18^x10-68^ Fisher’s exact test, p=1.00e^-3^ permutation test, see methods). Thus, the heterochromatinization of PODs in *Pogz^-/-^* cortex is accompanied by a collapse of TADs within these domains. Thus, the dramatic locus-level TAD collapse observed at the *Slitrk1,6,5* and *Lrfn5* example loci reflects a broader, POD-specific trend, with 61 of the altered TADs found in 34 PODs (Fig. 5E).

Because reduced intra-TAD interactions represented the dominant architectural change in *Pogz*⁻^/^⁻ cortex, we next asked whether TAD boundary integrity is also compromised at affected loci. TAD boundary integrity has been shown to localize clusters of regulatory elements within neighboring TADs closer together in 3D space to allow for proper transcriptional activity. Using cooltools to identify boundaries from Micro-C data (see Methods), we observed no global change in boundary strength across the genome (Fig. 5E). In contrast, TAD boundaries located within PODs exhibited a significant reduction in boundary strength in Pogz⁻^/^⁻ cortex (Fig. 5E). Overall, *Pogz* knockout mice display severe alterations in cortical TAD architecture and chromatin boundaries specifically within PODs.

### CTCF occupancy is reduced at weakened POD boundaries in *Pogz^⁻/⁻^* cortex

Next, we examined whether the architectural defects observed within PODs reflect disruption of the underlying boundary machinery. CCCTC-binding factor (CTCF) is a highly conserved zinc-finger protein that establishes many TAD boundaries and insulates adjacent chromatin domains (Phillips and Corces, 2009). We therefore asked whether CTCF occupancy is altered in *Pogz^⁻/⁻^* cortex. We performed CTCF ChIP-exo in E13.5 wildtype and *Pogz^-/-^* mouse cortex. ChIP-exo combines chromatin immunoprecipitation with exonuclease digestion to achieve near–base pair resolution of protein–DNA interactions (Rossi, Lai and Pugh, 2018; Rossi *et al*., 2021). Experiments were performed on pooled embryonic wildtype and *Pogz^-/-^* cortices (n = 5 per replicate). ChIP-exo data pileups demonstrate the base-pair resolution of binding at CTCF sequence motifs predicted genome-wide (Fig. S7A-D).

Genome-wide analysis identified 18,721 high-confidence CTCF binding sites in wildtype E13.5 cortex (Fig. S7I). Comparison of wildtype and *Pogz*⁻^/^⁻ samples revealed minimal differences in global CTCF occupancy, indicating that Pogz knockdown does not broadly alter CTCF binding across the genome (Fig. S7I). Globally, Micro-C signal piled up at CTCF binding sites and showed no changes in wildtype compared to the *Pogz^-/-^* (Fig. 6A). However, when concentrating on PODs, we observe that CTCF binding signal was markedly weakened in *Pogz^-/-^*mice (Fig. 6A). Similarly, CTCF binding signal at TAD boundaries was seemingly unchanged at a genome-wide scale but substantially reduced at boundaries that were lost within PODs by Micro-C (Fig. 6B,C). Thus, the collapse of TAD boundaries within PODs upon POGZ loss is accompanied by reduced CTCF binding.

**Figure 6.**
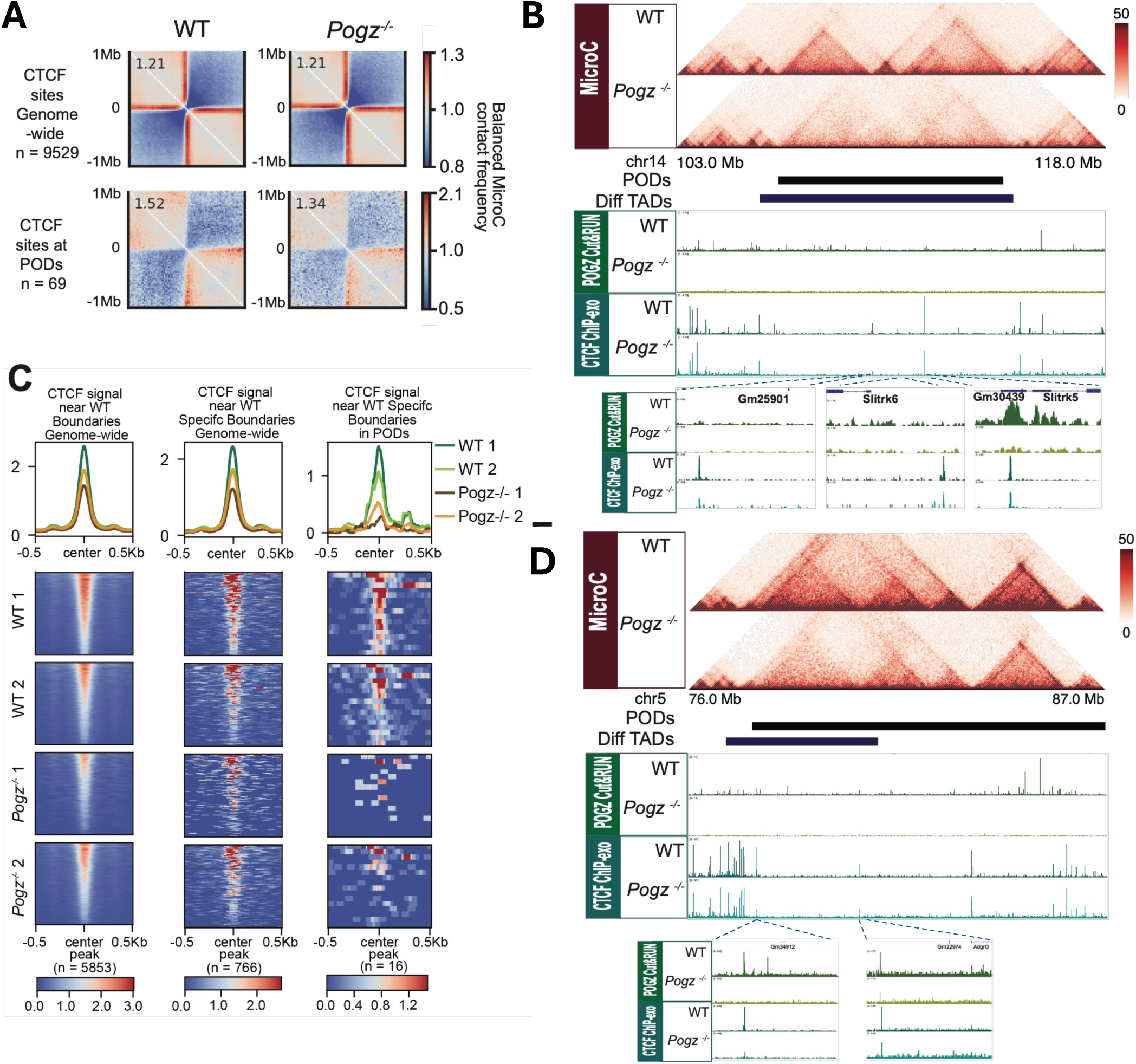
CTCF occupancy is reduced at weakened POD boundaries in *Pogz⁻ᐟ⁻* cortex. **(A)** Aggregate Micro-C contact frequency anchored at CTCF binding sites in wild-type and *Pogz⁻ᐟ⁻* cortex, genome-wide (n = 9,529) and at sites within PODs (n = 69). Corner values, mean enrichment. **(B)** Aggregate CTCF ChIP-exo signal (line plots and heatmaps; WT1, WT2, *Pogz⁻ᐟ⁻* 1, *Pogz⁻ᐟ⁻* 2) centered on wild-type TAD boundaries: genome-wide (n = 5,853), genome-wide subset (n = 766), and weakened boundaries within PODs (n = 16). **(C)** Locus view of the *Slitrk1/6/5* POD (chr14, 103.0–118.0 Mb). Tracks (top to bottom): Micro-C contact maps (WT, *Pogz⁻ᐟ⁻*); POD interval; differential TADs; CTCF ChIP-exo (WT, *Pogz⁻ᐟ⁻*). Inset: zoom at *Slitrk6/Slitrk5* highlighting co-localized POGZ CUT&RUN (WT and *Pogz⁻ᐟ⁻*) and CTCF ChIP-exo signal. **(D)** As in (C) for the *Adgrl3* POD (chr5, 76.0–87.0 Mb), with inset highlighting POGZ CUT&RUN (WT and *Pogz⁻ᐟ⁻*) and CTCF ChIP-exo at the locus.

To visualize how POD-associated CTCF loss manifests at individual genomic regions, we examined representative loci exhibiting robust heterochromatin gains and architectural disruption. At the *Slitrk* gene cluster and the *Adgrl3* locus, wildtype cortex displays well-defined TAD boundaries coincident with strong CTCF binding (Fig. 6B,D). In *Pogz*⁻^/^⁻, these same boundary elements show diminished CTCF ChIP-exo signal accompanied by local collapse or weakening of TAD organization, illustrating how POD-associated chromatin changes translate into focal architectural failure at neuronal gene loci. Notably, we observe co-localization of CTCF and POGZ CUT&RUN signal at altered boundaries near *Slitrk5* and *Adgrl3* (Fig. 6B,D).

### Heterochromatinization of PODs upon POGZ loss is accompanied by decreased transcription of neuronal genes harbored within PODs

To assess how Pogz impacts transcription during cortical development, we performed precision nuclear run-on sequencing (PRO-seq) on E13.5 wildtype and *Pogz^-/-^*mouse cortex. PRO-seq maps transcriptionally engaged RNA Polymerase II (PolII) at base-pair resolution, providing a direct readout of nascent RNA synthesis (Kwak *et al*., 2013). We generated biological replicate PRO-seq datasets from wild-type and *Pogz*⁻^/^⁻ cortex and quantified differential transcription using DESeq2 (see Methods).

Genome-wide, we identified 620 genes with significantly decreased nascent transcription and 469 genes with increased transcription in *Pogz*⁻^/^⁻ cortex (Fig. 7A; adjusted *p*-value < 0.05 and |log₂ fold change| > 0.585, see Methods). Gene ontology analysis revealed that down-regulated genes are strongly enriched for neurodevelopmental processes, including *forebrain development*, *generation of neurons*, *axon development*, and *regulation of cell communication*. In contrast, up-regulated genes were enriched for categories related to *nuclear pore organization* and *DNA damage response* (Fig. 7B, Fig. S8A). Consistent with these findings, POGZ has previously been implicated in neuronal development and DNA damage repair (Matsumura *et al*., 2020; Markenscoff-Papadimitriou *et al*., 2021; Li *et al*., 2025).

**Figure 7.**
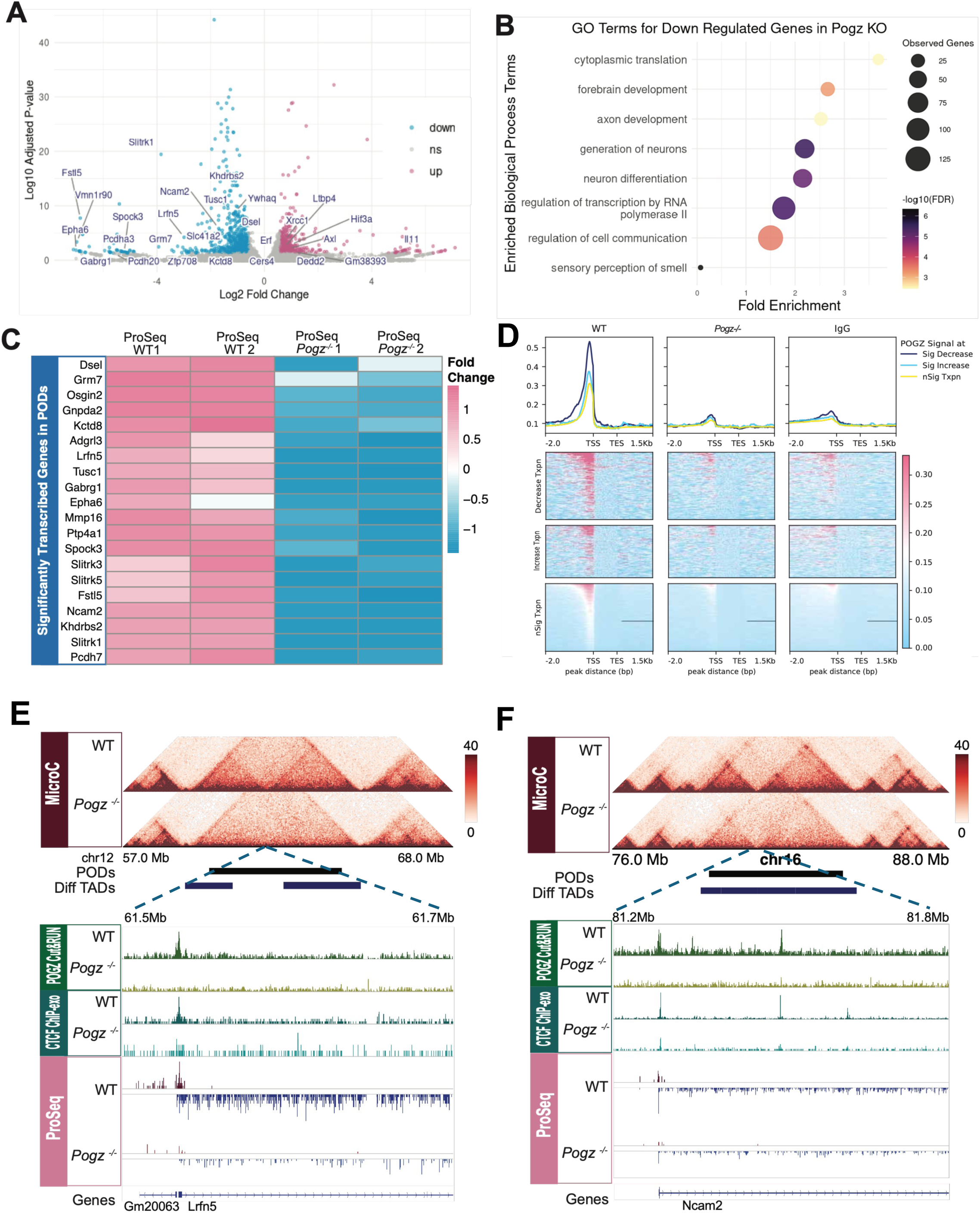
RNA Pol II nascent transcription is reduced at neurodevelopmental genes in *Pogz⁻ᐟ⁻*. (A) Volcano plot of differential nascent transcription (PRO-seq) in *Pogz⁻ᐟ⁻* versus wild-type E13.5 cortex. x-axis, log₂ fold change; y-axis, –log₁₀ adjusted *p*-value. Down-regulated genes (blue) and up-regulated genes (magenta) are colored; selected genes labeled. (B) GO Biological Process enrichment for down-regulated (top) and up-regulated (bottom) genes in *Pogz⁻ᐟ⁻*. Dot size, gene count; color, significance; x-axis, fold enrichment. (C) Heatmap of PRO-seq fold change for differentially transcribed genes within PODs across replicates (WT1, WT2, *Pogz⁻ᐟ⁻* 1, *Pogz⁻ᐟ⁻* 2). (D) Heatmaps of POGZ CUT&RUN signal in E13.5 wild-type cortex, centered on the transcription start sites (TSS) of significantly down-regulated (left) and up-regulated (right) genes in *Pogz⁻ᐟ⁻*. Each row is one gene; rows span −3.0 kb to +3.0 kb around the TSS. Color indicates enrichment score (es). (E) Locus view of the *Lrfn5* POD (chr12, 57.0–68.0 Mb). Tracks: Micro-C contact maps (WT, *Pogz⁻ᐟ⁻*); POD interval; differential TADs; and, at zoom POGZ CUT&RUN, CTCF ChIP-exo, and PRO-seq (each WT and *Pogz⁻ᐟ⁻*). RefSeq genes below. (F) As in (E) for the *Ncam2* POD (chr16).

To assess whether changes in nascent RNA levels are associated with POGZ binding, we examined the proximity of POGZ CUT&RUN sites to differentially transcribed genes. Genes with decreased nascent RNA in *Pogz*⁻^/^⁻ cortex were highly enriched near POGZ peaks (Fisher’s exact test, odds ratio = 2.63, p < 2.3 × 10⁻⁵; Fig. S8B). Strikingly, POGZ was found to occupy the promoters of essentially all down-regulated genes in *Pogz*⁻^/^⁻ mice (Fig. 7D), indicating that POGZ directly binds the genes that lose transcription upon its loss. Motif analysis of POGZ peaks at these promoters revealed strong enrichment for GC-rich Sp/KLF-family zinc-finger motifs, along with core-promoter (TATA) and Sox/HMG elements (Fig. S8C), consistent with POGZ occupying GC-box promoters of POGZ-dependent neuronal genes. These down-regulated genes were also slightly enriched within PODs (p = 0.033), whereas anti-POD regions showed no enrichment for differentially transcribed genes. Notably, all significantly differentially transcribed genes within PODs are down-regulated (Fig. 7C), consistent with increased heterochromatinization repressing transcription in these regions.

We illustrate these observations for several important neuronal genes. For example, the *Lrfn5* gene shows a strong reduction in nascent RNA levels in *Pogz*⁻^/^⁻ cortex (log₂FC = −2.99, adjusted log₁₀p = −4.21) and resides within a POD that contains a significantly altered topologically associating domain (TAD; p =0.05) as detected by Micro-C (Fig. 7E). Similarly, neural cell adhesion molecule 2 *(Ncam2)* is significantly down-regulated in *Pogz*⁻^/^⁻ cortex (log₂FC = −1.75, adjusted log₁₀ p = −6.35) and is located within a POD harboring a significantly differential TAD (p = 0.05) in *Pogz*⁻^/^⁻ Micro-C data (Fig. 7F). Notably, POGZ binds strongly at the promoters of both these genes, as does CTCF (Fig. 7E,F). Together, these results indicate that loss of POGZ leads to selective transcriptional repression of neuronal genes located within and outside of PODs.

### PODs converge on heterochromatin domains gained in Fragile X syndrome and POGZ shares binding sites with MeCP2

The selective heterochromatinization and silencing of long synaptic genes in *Pogz⁻^/^⁻* cortex closely resembles the biology of BREACHes, 36 megabase-scale heterochromatin domains in the human genome that silence synaptic genes in Fragile X syndrome cells carrying *FMR1* CGG expansions (Malachowski *et al*., 2023). To test whether these two phenomena affect a shared set of loci, we lifted over human BREACHes to the mouse genome (mBREACHes, see Methods) and compared their locations to the PODs. Of the 49 PODs, 6 directly overlapped mBREACHes (Fig. 8A), a notable overlap given that both domain classes are defined independently and arise from distinct genetic lesions in different species.

**Figure 8.**
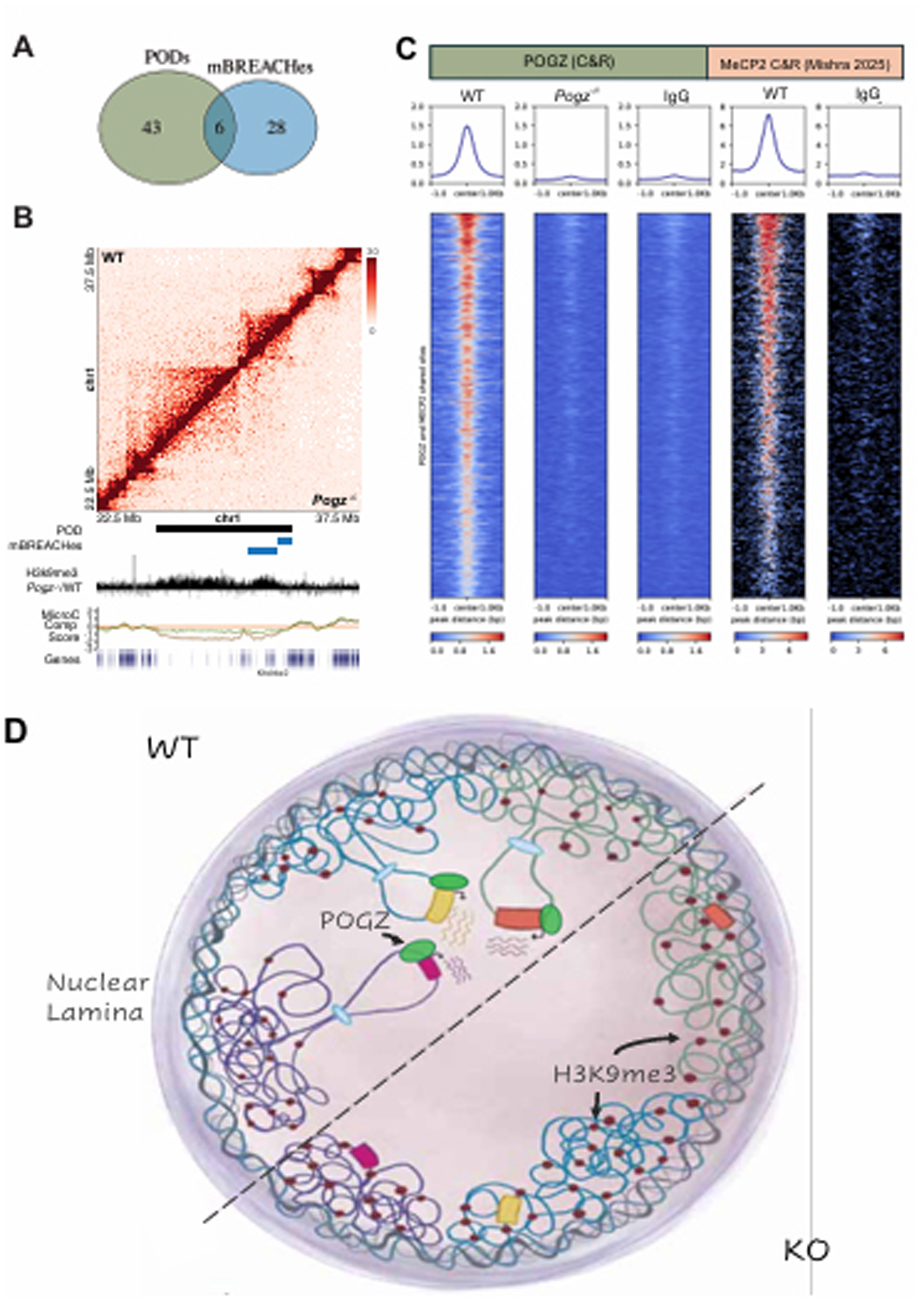
POGZ safeguards neuronal gene chromatin architecture, nuclear positioning, and transcription. **(A)** Venn diagram of overlap between PODs and mBREACHes (mouse orthologs of fragile X BREACHes) **(B)** Locus view of chr1 (22.5–37.5 Mb). Top: Micro-C contact map with wild-type (upper triangle) and *Pogz⁻ᐟ⁻* (lower triangle). Bottom tracks: POD and mBREACHes intervals H3K9me3 *Pogz⁻ᐟ⁻*/WT signal, Micro-C compartment score (WT in green and *Pogz⁻ᐟ⁻* in orange) and RefSeq genes (*Khdrbs2*). **(C)** Heatmaps and average profiles of POGZ CUT&RUN and MeCP2 CUT&RUN (Mishra et al., 2025) signal over POGZ/MeCP2 shared peaks (n = 1385) centered on peak ±1.0 kb and ranked by signal. **(D)** Schematic model of the cortical neuronal nucleus. In wild-type (top half), POD-resident neurodevelopmental gene loci reside in the nuclear interior in an active, looped configuration. In *Pogz⁻ᐟ⁻* (bottom half), these loci gain H3K9me3, strengthen B-compartment score, relocate to the nuclear lamina, and lose TAD/boundary architecture and transcription.

A representative shared locus spans the *Khdrbs2* region on chromosome 1, which is called as both a POD and an mBREACH (Fig. 8B). In *Pogz^-/-^* cortex, this domain gained H3K9me3, strengthened its B-compartment score by Micro-C, and lost local TAD structure relative to wild type mirroring the architectural collapse seen at *Slitrk*, *Lrfn5*, and *Adgrl3*, and recapitulating the heterochromatin spreading and 3D misfolding that defines BREACHes in Fragile X cells. PODs also converge on another independently-defined class of gene loci: several of the long synaptic genes they contain, including Slitrk1, Slitrk5, Lrfn5, Ncam2, and Flrt2, are “lonely genes,” neurodevelopmental cell-adhesion molecules isolated within massive, ancient gene deserts conserved across vertebrates (Chapman et al., 2026), linking PODs to a distinctive gene-desert architecture that is highly conserved.

A final line of convergence implicates a second NDD-associated regulator at these same loci. Comparing POGZ CUT&RUN from our cortical samples to MeCP2 CUT&RUN from adult mouse cortex (Mishra *et al*., 2025), we found that ∼20% of POGZ binding sites are co-occupied by MeCP2 (Fig. 8C). These co-bound sites are enriched for promoter-proximal peaks, and GREAT analysis of the shared regions returns GO terms centered on neurodevelopment such as “cerebral cortex neuron migration” and “Wnt signaling” and “cell cycle arrest” (Fig. S8D). Because MeCP2 is a core reader of methylated DNA at heterochromatin and is the disease gene in Rett syndrome, its presence at POGZ-bound loci in cortex suggests that POGZ may act within a broader gene regulatory network with MeCP2. Together with the mBREACH overlap, these results position POGZ at a convergence point for multiple NDD-linked regulators of chromatin, where distinct genetic lesions in POGZ, FMR1, and the MeCP2 pathway funnel onto common subsets of neuronal genes.

## Discussion

In this study, we identify POGZ as a binding partner of the H3K9 methyltransferase complex G9a/GLP and other heterochromatin factors in the developing mouse cortex, and show that POGZ loss drives bidirectional, megabase-scale redistribution of H3K9me3, accompanied by changes in 3D genome architecture and transcription at neurodevelopmental gene domains. Neuronal gene clusters that gain ectopic H3K9me3 in *Pogz^-/-^* cortex are repositioned to the nuclear lamina and exhibit strengthened B-compartment scores, locus-restricted erosion of TAD architecture, weakened boundary insulation, and reduced CTCF occupancy, with coordinated loss of transcription at neuronal genes (Fig. 8D). These findings establish POGZ as a G9a/GLP-associated chromatin regulator that maintains neuronal gene heterochromatin state, 3D chromatin architecture, and transcription during cortical development.

Together, these findings reveal that POGZ acts bidirectionally at the euchromatin–heterochromatin interface, promoting H3K9me3 at one set of neuronal loci while protecting another from it, a duality we develop below into a “sink-and-source” model. Notably, the domains most vulnerable to this dysregulation are synaptic gene clusters, the same class of genes targeted in Fragile X and other neurodevelopmental disorders, pointing to a potentially shared chromatin mechanism across distinct genetic lesions and common neurodevelopmental phenotypes.

### Heterochromatin regulation of neuronal gene expression

Our findings add to a growing appreciation that heterochromatin is not simply a default repressive state found at centromeres, telomeres, and repetitive elements, but an important regulator of neuronal gene expression. Since the discovery of H3K9me3 enrichment at olfactory receptor (OR) gene clusters in olfactory sensory neurons (OSNs)(Magklara *et al*., 2011), it has been increasingly appreciated that heterochromatin can promote neuronal gene expression in stochastic regulatory regimes. G9a-mediated H3K9 methylation is required for OR gene expression, and the H3K9 demethylase LSD1 is critical for productive transcription in this system (Lyons *et al*., 2013, 2014) (Lyons et al., 2013, 2014). Heterochromatin similarly organizes the clustered protocadherin (Pcdh) locus in olfactory neurons, where it may control variable isoform choice essential for neuronal wiring (Canzio *et al*., 2019). It is therefore striking that anti-PODs, the loci that lose H3K9me3 in *Pogz^-/-^* cortex, include OR and clustered protocadherin gene families, exactly the gene classes where H3K9me3 is known to be necessary and instructive for gene expression in olfactory neurons. Conversely, PODs, the loci that gain ectopic H3K9me3 in *Pogz*⁻^/^⁻ cortex, are enriched for large synaptic gene clusters (Slitrk, Adgrl3, Lrfn5, Ncam2, non-clustered Pcdh’s) embedded in gene-poor regions of the genome that are themselves moderately heterochromatic. Our findings suggest that POGZ protects these neuronal genes from becoming further heterochromatinized, and that this counter-silencing activity is required for their transcription. POGZ thus appears to perform two opposite functions at the interface of euchromatin and heterochromatin: facilitating G9a/GLP-mediated H3K9 methylation at gene clusters that require it in neurons, while shielding other neuronal gene domains from inappropriate spreading of the same mark.

### A sink-and-source model for POGZ function

We propose that POGZ acts as a context-dependent targeting factor for the G9a/GLP complex, with opposite consequences at different genomic locations. At anti-PODs (“source”), POGZ binding facilitates G9a/GLP recruitment and H3K9 methylation at gene clusters including ORs and protocadherins. At PODs (“sink”), POGZ binds open chromatin regions within neuronal gene domains encoding synaptic cell adhesion molecules, where it helps maintain accessibility and locally restrict heterochromatin spreading; its loss redistributes H3K9me3 into these domains. This logic has precedent in Polycomb biology, where loss of PRC2 anchors causes redistribution of H3K27me3, with depletion at normal targets and ectopic gains elsewhere (Fang *et al*., 2018; Harutyunyan *et al*., 2019; Kraft *et al*., 2022); as well as in KRAB-ZFP/KAP1 complexes that anchor H3K9me3 deposition at specific sites (Groner *et al*., 2010). We propose POGZ serves an analogous targeting role for G9a/GLP in the developing cortex, with bidirectional consequences that, when lost, affect heterochromatin across ∼10% of the genome.

### The many faces of POGZ

Most mechanistic studies of POGZ have relied on transformed human cell lines or pluripotent stem cells and have reached differing conclusions about its activity. In these systems, POGZ has been assigned to heterochromatin formation at pericentromeres and transposable elements through association with HP1, CHAMP1 and ADNP (Li *et al*., 2022, 2025), and, more recently, to a non-canonical Polycomb complex, PRC1.6, in which POGZ co-localizes with RING1B and HP1γ to repress BMP signaling during neuronal differentiation (Chavez *et al*., 2026). Indeed, in the developing cortex, POGZ co-occupies the genome with ADNP/ChAHP repressive complex at the majority of its binding sites, yet at the PODs and anti-PODs where it specifically governs H3K9me3 deposition, this co-occupancy is not observed, with POGZ binding these domains independently of ADNP (Fig. S6G). This separates POGZ’s canonical role in the ChAHP complex from its functions regulating heterochromatin at PODs and anti-PODs. Through the partner complex it engages at a given locus and developmental stage, POGZ produces bidirectional, locus-specific outcomes rather than acting as a dedicated silencer or activator.

### An evolutionary origin for POGZ counter-silencing

The ability of POGZ to oppose heterochromatin spreading at PODs may be a legacy of its evolutionary origin as a domesticated Pogo-family transposase (Cosby *et al*., 2021). Anti-heterochromatin activity is a recurrent theme among transposase-derived host proteins. In *Arabidopsis*, the Harbinger/PIF-superfamily-derived proteins ALP1 and ALP2 (Antagonist of Like Heterochromatin Protein 1 and 2) incorporate into PRC2 and antagonize Polycomb silencing, and rice encodes an apparently independently domesticated counterpart, PANDA, with comparable anti-silencing activity (Liang *et al*., 2015; Mao *et al*., 2022). Thus, multiple, independently domesticated transposases have converged on antagonizing heterochromatin, raising the possibility that POGZ’s counter-silencing function at PODs reflects an ancestral, transposon-derived capacity to keep host chromatin permissive, plausibly a trait that originally favored transposon expression and was later co-opted for neuronal gene regulation. This idea is speculative and remains to be tested, but it offers an evolutionary rationale for why a single transposase-derived protein would carry both silencing and counter-silencing activities. Intriguingly, the loci POGZ protects are themselves evolutionarily ancient: many POD genes are “lonely genes,” neurodevelopmental genes isolated within massive gene deserts highly conserved across vertebrates (Chapman *et al*., 2026). The pairing of an ancient, transposase-derived regulator with an ancient class of target genes suggests that the dependence of these vulnerable loci on specialized chromatin factors may be a deeply conserved feature of vertebrate genome regulation.

### A convergent mechanism across neurodevelopmental disorders

The chromatin architecture defects in Pogz^-/-^ cortex echo phenotypes in other genetically defined NDDs and point to a class of neuronal genes especially susceptible to heterochromatin dysregulation. PODs are dominated by synaptic genes and neuronal cell adhesion molecules (Slitrk’s, Adgrl3, Lrfn5, Ncam2, Pcdh’s) embedded in gene-poor regions. Whether the moderate heterochromatin state of these loci in wild-type brain is instructive for their expression remains unclear. But their unique vulnerability to aberrant H3K9me3 spreading and concomitant transcriptional loss in *Pogz^-/-^*cortex establishes that their expression is heterochromatin-regulated.

Strikingly, PODs overlap with BREACHes, the megabase-scale heterochromatin domains that form in Fragile X cells carrying CGG expansions in FMR1 (Malachowski *et al*., 2023) (Fig. 8). Like PODs, BREACHes engulf long synaptic genes, silencing them through coordinated heterochromatinization and 3D genome misfolding. MeCP2, the Rett syndrome gene, produces phenotypes that also resemble POGZ loss: it stabilizes expression of long neuronal genes whose dysregulation is a hallmark of Rett Syndrome (Moore *et al*., 2025), paralleling the POD silencing we observe. Consistent with a direct relationship, ∼20% of POGZ binding sites are co-occupied by MeCP2 in cortex (Mishra *et al*., 2025), enriched at promoters near neurodevelopmental genes (Fig. 8C). CHAMP1, a principal POGZ interactor in our IP-MS data, is mutated in CHAMP1-related intellectual disability syndrome (Nagai *et al*., 2022) and promotes heterochromatin assembly and homology-directed repair together with POGZ (Li *et al*., 2022, 2025).

Beyond these partnerships, our data recover the canonical ChAHP complex at genome scale. ADNP is mutated in Helsmoortel–Van der Aa syndrome, one of the most frequent single-gene causes of ASD (Satterstrom *et al*., 2020). POGZ co-occupies the majority of its binding sites with ADNP (Fig. S6G), consistent with prior work defining POGZ, ADNP, HP1 and CHD4 as core ChAHP components (Markenscoff-Papadimitriou *et al*., 2021). This concordance validates our IP-MS and CUT&RUN datasets against an established complex and confirms that the bulk of POGZ occupancy reflects its canonical role in the ChAHP complex. Critically, this ChAHP-bound majority is spatially distinct from the PODs and anti-PODs where POGZ regulates H3K9me3. At those domains, POGZ binds independently of ADNP, separating its ChAHP function from its G9a/GLP-associated, domain-scale heterochromatin regulating activity. POGZ, CHAMP1, MeCP2, and ADNP are thus all NDD risk genes converging on heterochromatin organization at partially overlapping neurodevelopmental gene loci. Together, these convergent partnerships suggest that distinct NDD mutations destabilize a shared network, producing comparable architectural and transcriptional defects despite different molecular entry points.

### Disentangling cause and effect at PODs

A central question raised by our data is the causal order of the events we observe. POGZ loss is accompanied by H3K9me3 gain, B-compartment strengthening, TAD erosion, reduced CTCF occupancy, lamina tethering, and transcriptional silencing at PODs, but our cross-sectional measurements cannot establish which change drives the others. At least three non-exclusive models are consistent with the data. In a *transcription-first* model, POGZ directly sustains transcription of POD genes, consistent with its binding at down-regulated promoters, and the loss of transcriptional activity permits heterochromatin spreading and architectural collapse. In an *architecture-first* model, POGZ primarily maintains CTCF-anchored boundaries and TAD insulation, whose failure allows H3K9me3 to spread across domains and silence transcription at genes within. In a *heterochromatin-first* model, POGZ chiefly sets the local balance of H3K9 methylation through its G9a/GLP association, and ectopic H3K9me3 gain is the upstream event that drives compartment change, lamina association, boundary loss, and transcriptional repression. Our data are compatible with all three: POGZ binds active promoters (favoring the first), co-localizes with CTCF at lost boundaries (favoring the second), and interacts with the H3K9 methylation machinery (favoring the third). Distinguishing these models will require perturbations with temporal resolution and directionality, which will convert the correlations reported here into a causal pathway.

### Lamina association and boundary loss as silencing routes

Two features of PODs in *Pogz^-/-^*cortex could repress POD genes independently of H3K9me3 gains: their repositioning to the nuclear lamina and the collapse of their internal TAD boundaries. The nuclear lamina is an instructive regulatory compartment in neural lineages, and lamina detachment can unlock genes for activation during neural differentiation (Peric-Hupkes *et al*., 2010). Nuclear lamin is required for silencing-by-repositioning in *Drosophila* neuroblasts (Kohwi *et al*., 2013), and detachment of long neuronal genes from the nuclear lamina precedes their activation in the embryonic mouse cortex (Rullens *et al*., 2025). The inappropriate lamina tethering of POD genes, themselves synaptic and neuronal migration genes, in the *Pogz^-/-^* cortex may therefore constrain their induction rather than simply reflect heterochromatinization. Similarly, weakened boundaries and reduced CTCF occupancy could disrupt the enhancer–promoter contacts these domains depend on to drive gene transcription. We thus favor a model in which H3K9me3 gain, perinuclear sequestration, and boundary loss act in concert to silence POD genes.

### Limitations of the study

Several questions remain open. Our IP-MS identifies G9a/GLP and CHAMP1 as principal POGZ interactors, but the domains mediating these contacts and whether POGZ modulates G9a/GLP catalytic activity, remain unknown. Mapping the POGZ–G9a/GLP interface, and how NDD-linked mutations disrupt it, will be key to understanding White–Sutton syndrome and POGZ-associated ASD. Second, causality is unresolved. Whether H3K9me3 gain, architectural collapse, nuclear repositioning, or transcriptional loss is upstream cannot be established from our cross-sectional data. Third, cell-type resolution is lacking. POGZ is expressed across excitatory neurons, inhibitory neurons, and progenitors with no apparent specificity, and E13.5 *Pogz⁻^/^⁻* cortex shows no overt shift in cell type composition. Still, cell-type-resolved analyses are needed to test whether POD and anti-POD phenotypes are uniform across lineages. Extending this to patient-derived neurons will clarify relevance to human disease.

## RESOURCE AVAILABILITY

### Lead contact

Eirene Markenscoff-Papadimitriou

### Materials availability

All newly generated reagents, mice, custom DNA FISH probes, and processed resources are available from the lead contact upon request.

### Data and code availability|

Sequencing data (H3K9me3 ChIP-seq, Micro-C, CTCF ChIP-exo, PRO-seq, and PRO-cap) generated in this study will be deposited at the NCBI Gene Expression Omnibus (GEO) and made publicly available prior to publication, with accession numbers listed here. Source data for imaging quantifications will be deposited at Zenodo and made publicly available prior to publication. Analysis scripts will be deposited at GitHub and made publicly available prior to publication.

## EXPERIMENTAL MODEL AND SUBJECT DETAILS

### Generation of the Pogz-HA knock-in mouse

A 3×HA epitope tag was inserted in-frame immediately before the endogenous Pogz stop codon by CRISPR-Cas9–mediated homology-directed repair. An sgRNA targeting the C-terminal coding region adjacent to the stop codon (protospacer: TGATGGAGATCTAGGTGCTG; PAM: AGG) was selected using CRISPOR (Concordet & Haeussler, 2018) based on predicted on-target efficiency and minimal off-target potential. A single-stranded oligodeoxynucleotide (ssODN) repair template (236 nt) was designed encoding three tandem HA epitopes (YPYDVPDYA × 3) preceded by a flexible GSAGSAAGSGEF linker, flanked by 57 bp and 62 bp homology arms matching the genomic sequence upstream and downstream of the stop codon. Insertion of the HA cassette disrupts the sgRNA recognition sequence, preventing re-cutting of the repaired allele. Cas9 protein, sgRNA, and ssODN were delivered by pronuclear injection into C57BL/6J zygotes. Founders were screened by PCR across the insertion site (primers POGZ-Fwd1: GCCTTCACCAGCTTTTTGAG; POGZ-Rev1: GAGTCAATGCTGGGAGTGGT; expected products 218 bp WT, 335 bp HA-tagged) followed by BtsI restriction digestion, which cleaves a site within the inserted linker sequence to distinguish tagged (∼91 + 244 bp) from wild-type (uncut) alleles. Correct in-frame insertion was confirmed by Sanger sequencing. Founders were backcrossed to C57BL/6J for at least two generations before experimental use.

### Homozygous *Pogz*⁻*^/^*⁻ Mouse strains

The *Pogz* mutant allele used in this study was the constitutive *Pogz* deletion allele generated by Markenscoff-Papadimitriou et al. (2021). Briefly, the allele was produced by CRISPR-Cas9 pronuclear injection using sgRNAs targeting exons 1 and 6, resulting in a ∼10 kb deletion that introduces a premature stop codon. Homozygous *Pogz*⁻/⁻ embryos are lethal perinatally, while heterozygous *Pogz*⁺/⁻ mice are viable and fertile. Wild-type littermates from heterozygous intercrosses served as controls. All mice were maintained on a C57BL/6 background, with the original founder outcrossed to C57BL/6 for ten generations.

### Animal husbandry and ethics

All animal procedures were approved by the Cornell University Institutional Animal Care and Use Committee (IACUC) and conducted in accordance with NIH guidelines. Mice were housed in social cages on a 12-hour light/dark cycle with ad libitum access to food and water, and were monitored daily. For timed pregnancies, noon on the day of vaginal plug detection was designated embryonic day 0.5. Animals of both sexes were used in all analyses.

### Embryo collection and sex

Timed pregnancies were established by housing *Pogz*⁺/⁻ males with *Pogz*⁺/⁻ females overnight, with noon of the day of vaginal plug detection designated as embryonic day (E) 0.5. Pregnant dams were euthanized at E13.5, and embryos were collected in ice-cold PBS. Embryos were genotyped by PCR, and tissue from individual embryos was processed separately to allow comparison of *Pogz*⁻/⁻, *Pogz*⁺/⁻, and wild-type littermates. Embryos of both sexes were used in all analyses, and sex was not determined prior to tissue collection.

### Cortical tissue dissection

Embryonic brains were removed from E13.5 embryos in ice-cold PBS and the telencephalon was isolated under a dissecting microscope. The cortex and basal ganglia (lateral and medial ganglionic eminences) were microdissected separately and immediately processed or flash-frozen depending on the downstream assay. For all genomic assays — RNA-seq, ATAC-seq, ChIP-seq, and CUT&RUN — tissue from individual embryos was processed separately to preserve genotype-specific biological replicates.

### Co-immunoprecipitation of POGZ-HA from embryonic mouse cortex

Cortices were dissected from E13.5 *Pogz-HA* knock-in embryos and from stage-matched wildtype (untagged) embryos as a negative control. For each condition, lysate from five pooled cortices was split into four technical IP replicates. Tissue was resuspended in 250 µL ice-cold Lysis Buffer (50 mM HEPES pH 7.5, 150 mM NaCl, 5 mM EDTA, 1% NP-40, 2 mM MgCl₂, 15% glycerol) freshly supplemented with Protease Inhibitor Cocktail (Roche) and PhosSTOP (Roche). Samples were rotated at 4 °C for 40 min, sonicated in a cup-horn sonicator for 1 min at 40% amplitude, and clarified by centrifugation at maximum speed for 30 min at 4 °C. Supernatants were transferred to LoBind tubes and protein concentration was measured by BCA assay. For each condition, input protein was normalized so that all four IP replicates received equal protein amounts.

Alpaca anti-HA single-domain antibody magnetic beads (Sigma, SAB5900004) were used for immunoprecipitation. Per IP replicate, 10 µL of bead slurry was washed twice with 1 mL Wash Buffer (Lysis Buffer without protease/phosphatase inhibitors), resuspended in Wash Buffer, and distributed evenly across the cleared lysates. Sample volumes were brought to a minimum of 250 µL to ensure proper mixing, and samples were rotated overnight (∼16–18 h) at 4 °C. The following day, beads were collected on a magnetic stand and washed 2–3× with 800 µL Wash Buffer, with sample transfer to a fresh LoBind tube on the final wash. Bound proteins were eluted in 100 µL Elution Buffer (100 mM HEPES pH 7.5, 1% SDS) at 95 °C for 8–10 min. Eluates were collected on a magnetic stand and stored at −80 °C.

### Mass Spectrometry Sample Preparation and Acquisition

IP eluates were reduced with 200 mM TCEP (Thermo Fisher Scientific, 77720) with room temperature incubation for 30 min, then alkylated with 375 mM iodoacetamide incubated for 30 min at room temperature and protected from light. The proteins were precipitated and washed 3 times with a precipitation solution (50% acetone, 49.9% ethanol, 0.1% acetic acid) to remove the SDS buffer. Protein pellets were resuspended using 8 M Urea, then diluted to 2 M Urea with NaCl/Tris solution (50 mM Tris pH 8.0, 150 mM NaCl). Peptide digestion of the samples was performed using 500 ng of mass-spectrometry grade Trypsin Gold (Promega, V5280) and nutated for 16 h at 37 °C, then quenched with equal volume of 4% formic acid. The samples were loaded and desalted on Evotips according to the manufacturer’s protocol (Evosep, EV2011), then analyzed with an Evosep One (Evosep) liquid chromatography system coupled to a timsTOF HT mass spectrometer (Bruker). Peptide separation was performed using the standard Evosep 60 SPD method on an 8 cm by 150 µm capillary column packed in-house with 1.9 µm C18 beads (Dr. Maisch, r119.aq.0001). Mass spectrometric analysis was competed using data-independent acquisition mode with parallel accumulation-serial fragmentation (dia-PASEF). A total of 34 mass windows with 25 Da width and 2 mobility windows each were acquired with a ramp time of 75 ms, covering a mass range from 350 to 1250 Da and a mobility range from 0.64 1/K0 [Vs/cm-2] to 1.37 1/K0 [Vs/cm-2]. Collision energy was increased as a function of ion mobility from 20 eV at 0.6 1/K0 [Vs/cm-2] to 59 eV at 1.6 1/K0 [Vs/cm-2]. Before running the samples, the elution voltage was calibrated for 1/K0 ratios in the timsControl software (Bruker) using three ions from ESI-L Tuning Mix (Agilent, G1969-85000) (m/z 622, 922, 1222).

### Western Blot

Total protein was extracted from flash frozen E13.5 liver using SDS lysis buffer (50 mM Tris pH 8.0, 10 mM EDTA, 1% SDS, Sigma-fast EDTA-free protease inhibitor #S8830) in a dounce homogenizer. Total protein was quantified using DC protein assay (BioRad #5000111). After boiling at 95°C for 10 minutes in 1X SDS sample buffer (60 mM Tris pH 8.0, 2% SDS, 10% glycerol, 0.02% Bromophenol Blue, 20 mM DTT (Thermo #D1532)), 40 ug of protein was loaded per lane. Blots were incubated overnight in primary antibodies (Sigma #11867423001 rat anti-HA 1:500; abcam #ab167408 rabbit anti-POGZ 1:1,000) and 2 hours in secondary antibodies (Santa Cruz #sc-2357 mouse anti-rabbit IgG HRP 1:3,000; CST #7077S goat anti-rat IgG HRP 1:2,000) before imaging using LumiGLO Reagent (CST #7003S) and LiCor Odyssey.

### POGZ and G9a Co-Immunoprecipitation

Cortices from 6 wild-type and 6 *Pogz*-HA heterozygous E13.5 embryos were pooled in 500 uL of lysis buffer (50mM HEPES (ph7.5), 150mM NaCl, 5mM EDTA, 1% NP-40, 2mM MgCl2, 5% Glycerol) with freshly added protease and phosphatase inhibitors (Pierce Protease Inhibitor tablets EDTA-free, Thermo #A32955; PhosSTOP, Sigma #4906845001). Protein was extracted by rotating at 4°C for 40 minutes, sonicating in a cup horn sonicator for 1 minute at 40% amplitude, and collecting the supernatant after spinning down for 30 minutes at 4°C. Total protein was quantified using DC protein assay (BioRad #5000111) and 400 ug of protein was input into each immunoprecipitation reaction, with 5% input set aside for analysis. Each sample rotated overnight with 25uL of Alpaca anti-HA beads (Sigma #SAB5900004) at 4°C. The next day, beads were washed with lysis buffer and protein was eluted in 30 uL of 2X SDS sample buffer (120 mM Tris pH 8.0, 4% SDS, 20% glycerol, 0.04% Bromophenol Blue, 40 mM DTT (Thermo #D1532)) by boiling samples at 95°C for 10 minutes. Samples were then loaded onto 4–15% Mini-PROTEAN® TGX Stain-Free™ Protein Gels (BioRad #4568083) along with the 5% input samples. Blots were incubated overnight in primary antibodies (Sigma #11867423001 rat anti-HA 1:500; abcam #ab185050 rabbit anti-G9a 1:1,000) and 1.5 hours in secondary antibodies (Santa Cruz #sc-2357 mouse anti-rabbit IgG HRP 1:3,000; CST #7077S goat anti-rat IgG HRP 1:2,000) before imaging using LumiGLO Reagent (CST #7003S) and LiCor Odyssey.

### Native H3K9me3 ChIP-seq

ChIP-seq with H3K9me3 antibody (see STAR methods) on native chromatin isolated from neuronal nuclei was performed as previously described (Magklara *et al*., 2011; Markenscoff-Papadimitriou *et al*., 2020). From each dissection of E13.5 cortex, intact nuclei were isolated by douncing twenty times with the loose pestle and a glass dounce implement in 0.5 mL Buffer 1 (300mM sucrose, 60mM KCl, 15mM NaCl, 15mM Tris-HCl, pH 7.5, 5mM MgCl2, 0.1mM EGTA, 1mM DTT, 1.1mM PMSF, EDTA-free Protease inhibitors). Then nuclei were lysed on ice for 10 minutes after adding 0.5 mL Buffer 2 (300mM sucrose, 60mM KCl, 15mM NaCl, 15mM Tris-HCl, pH 7.5, 5mM MgCl2, 0.1mM EGTA, 0.1% NP-40, 1mM DTT, 1.1mM PMSF, EDTA-free Protease inhibitors). Nuclei were counted using trypan blue and 500,000 nuclei were spun down at 7,000rpm for ten minutes at 4C. Nuclei were resuspended in 0.250mL MNase buffer (320mM sucrose, 50mM Tris-HCl, pH 7.5, 4mM MgCl2, 1mM CaCl2, 1.1mM PMSF) and incubated in a 37C water bath with 2 μl Micrococcal Nuclease enzyme (NEB) for eight minutes. Micrococcal Nuclease digestion was stopped by adding 10 μl 0.5M EDTA, and chromatin was spun down for 10 minutes 10,000rpm 4C. Soluble fraction “S1” supernatant was saved at 4C overnight, and “S2” fraction was dialyzed overnight in 250uL dialysis buffer at 4C (1mm Tris-HCl pH 7.5, 0.2mM EDTA, 0.1mM PMSF, 50mM Sodium Butyrate, Protease Inhibitors). Next day S1 and S2 fractions were combined, 50 μl were saved as input, and Chromatin immunoprecipitation was set up in ChIP buffer: 50mM Tris, pH 7.5, 10mM EDTA, 125 mM NaCl1, 0.1% Tween. 250mM Sodium Butyrate was supplemented for H3K27ac ChIPs. 1 μl of antibody was added to 1mL chromatin in ChIP buffer and incubated overnight at 4C rotating. Protein A and Protein G beads (10 μl for each ChIP) were blocked overnight in 700uL ChIP buffer, 20 uL yeast tRNA (20mg/mL), and 300uL BSA (10mg/mL). Beads were washed three times for five minutes on ice in Wash buffer 1 (50 mM Tris, pH 7.5, 10mM EDTA, 125mM NaCl, 0.1% Tween-20, with protease inhibitors and 5mM sodium butyrate) and three times in Wash buffer 2 (50 mM Tris, pH 7.5, 10mM EDTA, 175mM NaCl, 0.1% NP-40, with protease inhibitors and 5mM sodium butyrate), and ChIP DNA was eluted in elution buffer at 37C and purified by phenol chloroform extraction and ethanol precipitation. Sequencing libraries were made using Nugen Ovation Ultralow V2 kit and quantified by Agilent High Sensitivity DNA kit on the Agilent bioanalyzer.

### Micro-C

The cortex was dissected from wild-type and *Pogz*⁻/⁻ E13.5 mouse telencephalon and flash-frozen. Micro-C libraries were generated using the Dovetail® Micro-C Kit and Library Module for Illumina® (Cantata Bio) according to the manufacturer’s instructions. Briefly, ∼20 mg of tissue per sample was ground to a powder and dual-crosslinked with 3 mM disuccinimidyl glutarate (DSG) followed by ∼1% formaldehyde at room temperature. Crosslinked tissue was sequentially filtered through 200 μm and 50 μm filters to obtain a single-nucleus suspension, and chromatin was digested *in situ* with Micrococcal Nuclease (MNase) at 22°C for 15 minutes. Lysates were quantified by Qubit and digestion profiles confirmed to contain 40–70% mononucleosomes by Agilent Bioanalyzer. 1,000 ng of lysate was bound to Chromatin Capture Beads and processed through end polishing, bridge ligation with the biotinylated bridge oligonucleotide, and intra-aggregate ligation to generate proximity-ligated junctions marked with biotin. Crosslinks were reversed with Proteinase K, DNA was purified with SPRIselect beads (Beckman Coulter), and 150 ng was carried into library preparation comprising end repair, adapter ligation, and USER digest, without a fragmentation step. Adapter-ligated DNA was captured on Streptavidin Beads, washed, and amplified directly on-bead by 12 cycles of PCR with the Dovetail® Dual Index Primer Set #1 for Illumina®. A double-sided SPRIselect size selection was performed to recover fragments near the 350–1,000bp range. Libraries were sequenced on using Illumina PE150 with an average of 345 million reads per replicate.

### CUT&RUN

CUT&RUN was performed on freshly dissected E13.5 cortex from individual wild-type and *Pogz*⁻/⁻ embryos, with each embryo processed as a separate biological replicate. Cortices were kept in ice-cold Earle’s Balanced Salt Solution (EBSS) during embryo genotyping. Nuclei were isolated using Buffer 1 and Buffer 2 as described above for native H3K9me3 ChIP-seq, with permeabilization by ten strokes of a loose-fit Dounce pestle. Nuclei were pelleted at 600 × g and resuspended in Wash Buffer (20 mM HEPES pH 7.5, 150 mM NaCl, 0.5 mM spermidine, 1× EDTA-free protease inhibitor) from the EpiCypher CUTANA CUT&RUN protocol (v1.6, 8.4.2020).

Subsequent steps followed the EpiCypher CUTANA CUT&RUN protocol. Nuclei were bound to activated Concanavalin A magnetic beads, resuspended in cold Antibody Buffer (Wash Buffer + 0.01% digitonin + 2 mM EDTA), and incubated overnight at 4°C with target-specific antibody. The following day, beads were washed in Digitonin Buffer (Wash Buffer + 0.01% digitonin), incubated with pAG-MNase (CUTANA, EpiCypher) for 10 minutes at room temperature, washed again in Digitonin Buffer, and chromatin digestion was activated by addition of 2 mM CaCl₂ and incubation on a nutator for 2 hours at 4°C. Reactions were quenched with Stop Buffer (340 mM NaCl, 20 mM EDTA, 4 mM EGTA, 50 μg/mL RNase A, 50 μg/mL glycogen) and incubated at 37°C for 10 minutes to release fragments and degrade RNA. Released DNA was purified from the supernatant using the CUTANA DNA Purification Kit (EpiCypher) and quantified by Qubit dsDNA HS assay.

Sequencing libraries were prepared from CUT&RUN-enriched DNA using the NEBNext Ultra II DNA Library Prep Kit for Illumina (NEB E7645). Briefly, end repair was performed at 20°C for 30 minutes followed by 50°C for 1 hour to preserve small (25–70 bp) fragments characteristic of transcription factor footprints. Diluted NEBNext adapters (1:10) were ligated, USER-treated, and adapter-ligated DNA was cleaned up with 1.75× AMPure XP beads. Libraries were amplified by 17 cycles of PCR (98°C 10 s denaturation, 65°C 10 s combined annealing/extension) with NEBNext indexing primers, and double-sided AMPure XP size selection (0.8× then 1.2× final) was performed to remove fragments >350 bp and <150 bp. Final library size distribution was assessed on an Agilent TapeStation, and libraries were sequenced on an Illumina platform in paired-end mode.

### CTCF ChIP-exo

CTCF ChIP-exo was performed on E13.5 wild-type and *Pogz*⁻/⁻ mouse cortex following the ChIP-exo 5.0 protocol (Rossi et al., 2018; Rossi et al., 2021). Briefly, dissected cortex was crosslinked in 1% formaldehyde, quenched with 125 mM glycine, and washed. Cells were lysed and chromatin was solubilized and sonicated to fragments of ∼100–500 bp. Sonicated chromatin was immunoprecipitated overnight at 4°C with an anti-CTCF antibody (Millipore 07-729) bound to Protein G Dynabeads. Bead-bound chromatin was processed through the streamlined on-bead enzymatic series of ChIP-exo 5.0: A-tailing with Klenow fragment (3′→5′ exo⁻); combined T4 polynucleotide kinase and T4 DNA ligase reaction in PEG-containing buffer to attach the first sequencing adapter; and lambda exonuclease digestion to trim the 5′ ends of DNA up to the protein–DNA crosslink, generating a near base-pair-resolution boundary. Following crosslink reversal with proteinase K and DNA purification, the second adapter was attached by splint-mediated ssDNA ligation to the resected 3′ end. Libraries were PCR-amplified, size-selected on an agarose gel to recover fragments of 200–500 bp, and sequenced on an Illumina platform in paired-end mode.

### PRO-seq

Cells from individual E13.5 cortices were permeabilized in buffer containing 10 mM Tris-HCl pH 7.4, 10 mM KCl, 300 mM sucrose, 5 mM MgCl₂, 1 mM EGTA, 0.1% NP-40, 0.05% Tween-20, 0.5 mM DTT, protease inhibitors, and SUPERase-In, with a 2% spike-in of *Drosophila* S2 cells, and resuspended in storage buffer (10 mM Tris-HCl pH 8.0, 25% glycerol, 5 mM MgCl₂, 0.1 mM EDTA, 5 mM DTT). A two-biotin nuclear run-on was performed at 37°C for 5 minutes in run-on master mix (final concentrations: 10 mM Tris-HCl pH 8.0, 0.5% Sarkosyl, 5 mM MgCl₂, 300 mM KCl, 1 mM DTT, 40 μM each of ATP, GTP, Biotin-11-CTP, and Biotin-11-UTP, plus SUPERase-In). The reaction was stopped by addition of TRIzol LS for immediate RNA extraction. Total RNA was base-hydrolyzed with 0.2 N NaOH on ice for 10 minutes, neutralized with 1 M Tris-HCl pH 6.8, and buffer-exchanged through a Bio-Rad P30 column. Biotinylated nascent RNA was enriched on Streptavidin C1 magnetic beads (Thermo Fisher), eluted with TRIzol, and a 3′ adaptor (REV3+6N) was ligated with high-concentration T4 RNA Ligase I (T4 RNA Ligase I, HC) in the presence of 10% PEG-8000 at 4°C overnight on an end-to-end rotator. Following a second streptavidin capture, on-bead 5′ decapping was performed with RppH (NEB) at 37°C for 1 hour, followed by 5′ hydroxyl repair with T4 PNK at 37°C for 1 hour. A 5′ adaptor (Rev5+6N) was ligated on-bead with high-concentration T4 RNA Ligase I (HC) at 4°C overnight on an end-to-end rotator. RNA was eluted with TRIzol, reverse-transcribed with Maxima H-minus reverse transcriptase using primer RP1, and amplified with primer RP1 and barcoded RPI-n primers (cycle number determined by a test dilution series, typically ∼21 total cycles). The RP1 and RPI-n primer sequences were derived from the Illumina TruSeq Small RNA library prep kit and custom-synthesized by IDT. Libraries were size-selected from 170–300 bp on 8% native TBE PAGE gels to remove adaptor dimers, and sequenced paired-end 150 by Novogene on an Illumina NovaSeq X Plus.

### DNA Fluorescent In Situ Hybridization (FISH)

DNA FISH was performed as previously described (Markenscoff-Papadimitriou et al *Cell* 2014). Embryonic brains were dissected into cold PBS and fixed in 4% paraformaldehyde (PFA) at 4°C overnight, followed by three washes in cold PBS. Tissue was cryoprotected in 30% sucrose overnight at 4°C, then embedded in optimal cutting temperature (OCT) compound and flash-frozen on a liquid-nitrogen-cooled surface. Blocks were stored at −20°C until sectioning. Cryosections of 8 µm were collected onto Superfrost Plus slides, air-dried overnight, and stored at −20°C.

BAC DNA was isolated from BAC bacteria (CHORI BacPac) using the Nucleobond BAC prep kit (see Supplementary Table). Approximately 1 µg of BAC DNA was labeled by nick translation in a 20 µL reaction containing 4 µL of biotin-or digoxigenin (DIG)-conjugated nick translation mix, incubated at 15°C for 3 hours. Reactions were quenched by addition of 0.5 M EDTA (1 µL) and heat inactivation at 65°C for 10 minutes, then purified by spin column. Probe fragment size was verified by gel electrophoresis to be 100-200 base pairs. Labeled probe was ethanol-precipitated with approximately 10× mass Cot-1 DNA and 50× mass salmon sperm DNA to suppress repetitive sequence hybridization, air-dried, and resuspended in hybridization buffer (50% formamide, 2× SSC, 0.02% Denhardt’s solution, 1% dextran sulfate).

Slides were warmed to room temperature in a closed box for 1–2 hours to prevent condensation. Heat-induced antigen retrieval was performed by immersing slides in 10 mM sodium citrate (pH 6.0) heated just below simmering (approximately 95°C) for 10 minutes in a vegetable steamer, followed by immediate dehydration through a cold ethanol series (70% for 2 minutes, 90% for 2 minutes, 100% for 5 minutes, all on ice) and air-dried for 5 minutes on a slide warmer.

Probes were denatured at 75°C for 10 minutes and held at 37°C until use. Slides were denatured in 70% formamide / 2× SSC pre-warmed to 75°C for several hours, immersing slides for 10 minutes at 75°C. Slides were then re-dehydrated through the cold ethanol series described above and air-dried on a slide warmer. Approximately 5 µL of undiluted probe was applied per section, coverslipped with a 12mm circular glass coverslip, and sealed with rubber cement (Best-Test brand). Hybridization was carried out at 37°C for 12–18 hours

Rubber cement was removed under warm wash buffer (50% formamide / 2× SSC, approximately 45°C), and slides were washed twice for 10 minutes at 37°C, followed by two 5-minute washes in 0.1% Triton X-100/PBS at room temperature. Probe detection was performed overnight at 4°C using mouse anti-DIG (clone 499, 1:200) and rabbit anti-streptavidin conjugated to a 597 nm fluorophore (1:200) diluted in 0.1% Triton X-100/PBS. Slides were washed three times for 5 minutes, then mounted with DAPI-containing anti-fade mountant and coverslipped for imaging.

For combined immunofluorescence and DNA-FISH, the FISH protocol above was followed through the detection washes, after which slides were post-fixed in 4% PFA for 2 minutes at room temperature and washed twice in PBS/0.1% Triton X-100. Slides were blocked for 1 hour at room temperature in PBS/0.15% Triton X-100/4% donkey serum (PTS), protected from light. Primary antibodies targeting lamin B1 (1:200 in PTS) were applied overnight at 4°C. Slides were washed twice in PBS/0.1% Triton X-100, then incubated with fluorophore-conjugated secondary antibody (1:200 in PTS, 2 µL per 400 µL) for 2 hours at room temperature. Following two additional washes in PBS/0.1% Triton X-100, slides were mounted with DAPI-containing mountant and imaged.

### Microscopy

Slides were imaged on a Leica Stellaris 3 confocal microscope using the Leica Application Suite X software. Images were acquired at 63× magnification as tiled z-stacks spanning the full thickness of each section, and tiles were stitched in LAS X to generate full-section images at single-cell resolution. Acquisition parameters (Smart Gain and Smart Intensity) were held constant across all sections and genotypes within an experiment to allow quantitative comparison: DAPI for nuclei, Alexa Fluor 546 for lamin B1 immunostaining, and Alexa Fluor 488 for DNA FISH puncta. Image processing was likewise standardized across wild-type and *Pogz*⁻/⁻ samples: each channel was opened in Fiji, adjusted with auto-brightness/contrast, assigned a consistent LUT (blue for DAPI, magenta for lamin B1, green for DNA FISH), converted to 16-bit.

### Lamina contact analysis

Image processing for DNA FISH and Lamin B1 co-localization was performed as in Begnis et al. (2024), with manual scoring of lamina contacts. Following acquisition of z-stacks, images were processed in Fiji. A nucleus was scored as having a lamina contact at a given locus when ≥1 DNA FISH punctum (Alexa Fluor 488, green) directly overlapped with the lamin B1 immunostaining signal (Alexa Fluor 546, red), producing a yellow signal in the merged image. Each nucleus was scored independently across all visible alleles, and the proportion of nuclei with ≥1 colocalized punctum was reported per embryo. All scoring was performed manually by a single observer to ensure consistency, and embryos were used as biological replicates. Statistical comparisons between genotypes were performed with two-tailed Welch’s t-tests (unequal variance assumed).

## QUANTIFICATION AND STATISTICAL ANALYSIS

### IP-MS analysis

Raw mass spectrometry data acquired in data-independent acquisition (DIA) mode were processed using DIA-NN (version 2.0). Spectra were searched against the reviewed mouse UniProtKB/Swiss-Prot protein database (release January 2024; taxonomy ID: 10090) using DIA-NN in library-free mode. Database searching allowed up to three missed tryptic cleavages and a maximum of two variable modifications per peptide. Carbamidomethylation of cysteine residues was specified as a fixed modification, while methionine oxidation was included as a variable modification. N-terminal methionine excision was enabled during the search. The fragment ion m/z range was set to 100–1800. Match-between-runs (MBR) was enabled, and peptide quantification was performed using the QuantUMS (high precision) strategy (Grossmann *et al*., 2026). Peptide and protein identifications were filtered at a 1% false discovery rate (FDR) using the default DIA-NN settings (Demichev *et al*., 2020).

DIA-NN output files were further processed and visualized using Python and R. Protein enrichment was evaluated by comparing four biological replicates of Pogz-HA immunoprecipitation with four biological replicates of the Pogz-WT control. For each protein, abundance ratios were calculated between four pairwise combinations of Pogz-HA and Pogz-WT samples, and the protein fold change (FC) was defined as the median of these pairwise abundance ratios. Statistical significance (raw P values) was assessed using an empirical Bayes linear modeling framework implemented in the limma package (Ritchie *et al*., 2015). This approach moderates variance estimates by borrowing information across all quantified proteins, thereby improving statistical power and reducing false-positive discoveries arising from noisy measurements. Multiple-testing correction was performed using the Benjamini–Hochberg procedure to control the false discovery rate (FDR), yielding adjusted P values. Proteins were considered significant Pogz interactors if they met the criteria of log2FC > 1 and adjusted P < 0.05.

### H3K9me3 ChIP-seq analysis

Paired-end H3K9me3 ChIP-seq reads from wild-type, and knockout samples were quality checked and adapter-trimmed using Trim Galore (paired mode) with a stringency of 4, a minimum read length of 20 bp, and concurrently assessed with FastQC. Trimmed reads were aligned to the mouse reference genome (mm10) using Bowtie2. The resulting alignments were sorted, and duplicates were marked with Picard (v2.26.1) and removed to retain only non-duplicate reads. Individual H3K9me3-enriched regions were called from the deduplicated files using SICER, with the matched sequencing input serving as the control. Given the broad distribution characteristic of H3K9me3, peaks were identified against the mm10 genome using a window size of 300 bp, a gap size of 3000 bp, a fragment size of 1000 bp, and a false discovery rate (FDR) threshold of 0.01. To identify PODs and AntiPODs, the same parameters were applied to differential enrichment analyses comparing WT versus KO and KO versus WT. Results with KO as treatment and WT as control resulted in bed regions that tended to cluster together. To capture these enriched sites, BED regions from replicate results were merged: adjacent regions within 2 Mb were combined into single regions, and the resulting regions shorter than 500 kb were discarded. This same process was then used to capture broad megabase scale regions with decreased H3K9me3 in KO.

### Micro-C processing and quality control

Micro-C data were processed using the distiller-nf pipeline (Open2C). Paired-end reads from each library were aligned to the mouse reference genome (mm10) using BWA-MEM. Reads were parsed into Hi-C pairs and optical duplicates removed with pairtools. A maximum mismatch of 1 bp between mapped positions on either side for two pairs to be considered duplicates. Deduplicated pairs for individual and merges libraries were binned into contact matrices using cooler and stored as multi-resolution.mcool files spanning resolutions from 1 kb to 10 Mb. All contact matrices were iteratively balanced normalized by iterative correction (IC; Imakaev et al. 2012) as implemented in cooler (Abdennur and Mirny 2020) and used for downstream analysis.

### Compartment analysis

A/B compartments were identified by eigenvector decomposition using the eigs-cis function of cooltools (Open2C). For each individual and each merged libraries, intra-chromosomal eigenvectors were computed at 100 kb resolution cool files. H3K27ac ChIP-seq signal track binned at 100 kb was used to orient the first eigenvector (E1) and assign A or B compartments. Differential compartmentalization between WT and KO was assessed across autosomes, after removing low-coverage or low-confidence bins. Concordant compartment scores were evaluated by Pearson correlation between WT and KO E1 values generated by cooltools eigs-cis and overall compartment change were tested with a paired t-test. To identify bins with a significant change in compartment strength score, the E1 difference (KO − WT) was compared against the genome-wide difference distribution. P-values were corrected for multiple testing using the Benjamini–Hochberg false discovery rate (FDR), and bins with an adjusted p < 0.1 were considered significant. Significant bins were further classified by the sign of their E1 values and the direction of compartment change.

### TAD and boundary analysis

Insulation scores and boundaries were called using 25 kb-resolution contact matrices for both WT and KO libraries (for both individual and merged replicates) using the insulation function of cooltools (Open2C et al. 2024). Insulation profiles were computed across a range of sliding diamond-window, using the iterative-correction balancing weights Boundary strength was thresholded using the Li method. TADs called for the WT and KO was done using the hicFindTADs function from HiCExplorer (v3.7.2; Ramírez et al. 2018). Differential TADs were then identified with hicDifferentialTAD, using WT TADs as the reference set compared against the matched KO contact matrices. Differences were assessed with a Wilcoxon rank-sum test, rejecting the null hypothesis that the two conditions are equal at a p-value threshold of ≤ 0.05. Under the “all” rejection mode, a TAD was classified as differential only when all tested regions (the intra-TAD region and both inter-TAD regions) showed a significant difference, yielding a set of high-confidence differential TADs between WT and KO.

### CTCF ChIP-exo analysis

Raw paired-end sequencing reads were trimmed with Trim Galore using a stringency of 4, a minimum read length of 20 bp, and paired-end mode, with post-trimming quality assessed by FastQC. Trimmed reads were aligned to the mouse reference genome (mm10) using Bowtie2 with default parameters. Aligned reads were filtered with SAMtools to retain only properly paired reads (-f 2) with a minimum mapping quality of 10, and reads mapping to chrM, chrY, and unplaced or random contigs were removed. PCR and optical duplicates were then marked and removed using Picard MarkDuplicates (v2.19.2) with REMOVE_DUPLICATES=true and coordinate-sorted input. Peaks were called from the deduplicated BAM files against matched IgG controls using MACS3 callpeak with the mouse genome size (-g mm), broad peak detection (--broad-cutoff 0.05), retention of all remaining reads (--keep-dup all), signal-per-million-reads normalization (--SPMR), bedGraph output (-B), and model building disabled in favor of a fixed 75 bp fragment extension (--nomodel --extsize 75) to accommodate the exonuclease-trimmed footprints characteristic of the CTCF ChIP-exo protocol.

### PRO-seq analysis

Raw paired-end PRO-seq reads were processed using the proseq2.0 pipeline (Chu, T., Wang, Z., Chou, S. P., & Danko, C. G. (2018). Discovering Transcriptional Regulatory Elements From Run-On and Sequencing Data Using the Web-Based dREG Gateway. Current protocols in bioinformatics, e70.). Unique molecular identifiers (UMIs; 6 nt) were trimmed from both read ends, and adapter sequences were removed using cutadapt. PCR duplicates were eliminated by UMI-based deduplication. Reads were aligned to the mouse reference genome (mm10) using BWA, and read quality was assessed with FastQC. Gene-level quantification was performed with featureCounts (v2.0.6) in reverse-strand mode, using GENCODE mm10 gene annotations. Differential expression analysis was carried out with DESeq2; genes with an adjusted *p*-value < 0.05 and |log₂ fold change| > 0.585 were considered differentially expressed.

## Notes

### Competing Interest Statement

The authors have declared no competing interest.

### Funding

- Simons Foundation: SFARI893926, SFARI868055

## References

Abelson, J.F. et al. (2005) “Sequence variants in SLITRK1 are associated with Tourette’s syndrome,” Science (New York, N.Y.), 310(5746), pp. 317–320.

Batzir, N.A. et al. (2020) “Phenotypic expansion of POGZ-related intellectual disability syndrome (White-Sutton syndrome),” American Journal of Medical Genetics Part A, pp. 38–52. Available at: 10.1002/ajmg.a.61380.

Canzio, D. et al. (2019) “Antisense lncRNA Transcription Mediates DNA Demethylation to Drive Stochastic Protocadherin α Promoter Choice,” Cell, pp. 639–653.e15. Available at: 10.1016/j.cell.2019.03.008.

Chapman, M.A. et al. (2026) “Human neurodevelopmental genes housed in massive, ancient gene deserts,” bioRxiv. bioRxiv. Available at: 10.64898/2026.03.27.714728.

Chavez, J. et al. (2026) “The zinc-finger protein POGZ associates with polycomb Repressive Complex 1 to regulate bone morphogenetic protein signaling during neuronal differentiation,” Stem cell reviews and reports, 22(3), pp. 1371–1387.

Cosby, R.L. et al. (2021) “Recurrent evolution of vertebrate transcription factors by transposase capture,” Science, 371(6531). Available at: 10.1126/science.abc6405.

Cunniff, M.M. et al. (2020) “Altered hippocampal-prefrontal communication during anxiety-related avoidance in mice deficient for the autism-associated gene Pogz,” eLife, 9. Available at: 10.7554/eLife.54835.

De Rubeis, S. et al. (2014) “Synaptic, transcriptional and chromatin genes disrupted in autism,” Nature, 515(7526), pp. 209–215.

Deciphering Developmental Disorders Study (2015) “Large-scale discovery of novel genetic causes of developmental disorders,” Nature, 519(7542), pp. 223–228.

Demichev, V. et al. (2020) “DIA-NN: neural networks and interference correction enable deep proteome coverage in high throughput,” Nature Methods, 17(1), pp. 41–44.

Fang, D. et al. (2018) “H3.3K27M mutant proteins reprogram epigenome by sequestering the PRC2 complex to poised enhancers,” eLife, 7, p. e36696.

Groner, A.C. et al. (2010) “KRAB-zinc finger proteins and KAP1 can mediate long-range transcriptional repression through heterochromatin spreading,” PLoS Genetics, 6(3), p. e1000869.

Grossmann, J.L. et al. (2026) “Accurate quantification in proteomics with QuantUMS,” Nature Biotechnology [Preprint]. Available at: 10.1038/s41587-026-03131-2.

Harutyunyan, A.S. et al. (2019) “H3K27M induces defective chromatin spread of PRC2-mediated repressive H3K27me2/me3 and is essential for glioma tumorigenesis,” Nature Communications, 10(1), p. 1262.

Heath, J. et al. (2022) “POGZ promotes homology-directed DNA repair in an HP1-dependent manner,” EMBO reports, 23(1), p. e51041.

Iossifov, I. et al. (2014) “The contribution of de novo coding mutations to autism spectrum disorder,” Nature, 515(7526), pp. 216–221.

Kiefer, L. et al. (2023) “WAPL functions as a rheostat of Protocadherin isoform diversity that controls neural wiring,” *Science (New York*, N.Y.), 380(6651), p. eadf8440.

Kohwi, M. et al. (2013) “Developmentally regulated subnuclear genome reorganization restricts neural progenitor competence in Drosophila,” Cell, 152(1–2), pp. 97–108.

Kraft, K. et al. (2022) “Polycomb-mediated genome architecture enables long-range spreading of H3K27 methylation,” Proceedings of the National Academy of Sciences of the United States of America, 119(22), p. e2201883119.

Kwak, H. et al. (2013) “Precise maps of RNA polymerase reveal how promoters direct initiation and pausing,” Science, 339(6122), pp. 950–953.

Li, F. et al. (2022) “CHAMP1 binds to REV7/FANCV and promotes homologous recombination repair,” Cell reports, 40(9), p. 111297.

Li, F. et al. (2025) “CHAMP1 complex directs heterochromatin assembly and promotes homology-directed DNA repair,” Nature communications, 16(1), p. 1714.

Liang, S.C. et al. (2015) “Kicking against the PRCs - A domesticated transposase antagonises silencing mediated by Polycomb group proteins and is an accessory component of Polycomb repressive complex 2,” PLoS Genetics, 11(12), p. e1005660.

Lieberman-Aiden, E. et al. (2009) “Comprehensive mapping of long-range interactions reveals folding principles of the human genome,” *Science (New York*, N.Y.), 326(5950), pp. 289–293.

Lyons, D.B. et al. (2013) “An epigenetic trap stabilizes singular olfactory receptor expression,” Cell, 154(2), pp. 325–336.

Lyons, D.B. et al. (2014) “Heterochromatin-mediated gene silencing facilitates the diversification of olfactory neurons,” Cell Reports, 9(3), pp. 884–892.

Magklara, A. et al. (2011) “An epigenetic signature for monoallelic olfactory receptor expression,” Cell, 145(4), pp. 555–570.

Malachowski, T. et al. (2023) “Spatially coordinated heterochromatinization of long synaptic genes in fragile X syndrome,” Cell, 186(26), pp. 5840-5858.e36.

Mao, D. et al. (2022) “The Harbinger transposon-derived gene PANDA epigenetically coordinates panicle number and grain size in rice,” Plant Biotechnology Journal, 20(6), pp. 1154–1166.

Markenscoff-Papadimitriou, E. et al. (2020) “A Chromatin Accessibility Atlas of the Developing Human Telencephalon,” Cell, 182(3), pp. 754–769.e18.

Markenscoff-Papadimitriou, E. et al. (2021) “Autism risk gene POGZ promotes chromatin accessibility and expression of clustered synaptic genes,” Cell reports, 37(10), p. 110089.

Matsumura, K. et al. (2020) “Pathogenic POGZ mutation causes impaired cortical development and reversible autism-like phenotypes,” Nature communications, 11(1), p. 859.

McLean, C.Y. et al. (2010) “GREAT improves functional interpretation of cis-regulatory regions,” Nature biotechnology, 28(5), pp. 495–501.

Mishra, G.P. et al. (2025) “Interaction of methyl-CpG-binding protein 2 (MeCP2) with distinct enhancers in the mouse cortex,” Nature Neuroscience, 28(1), pp. 62–71.

Moore, J.R. et al. (2025) “MeCP2 and non-CG DNA methylation stabilize the expression of long genes that distinguish closely related neuron types,” Nature Neuroscience, 28(6), pp. 1185–1198.

Nagai, M. et al. (2022) “Deficiency of CHAMP1, a gene related to intellectual disability, causes impaired neuronal development and a mild behavioural phenotype,” Brain communications, 4(5), p. fcac220.

Nozawa, R.-S. et al. (2010) “Human POGZ modulates dissociation of HP1α from mitotic chromosome arms through Aurora B activation,” Nature cell biology, 12(7), pp. 719–727.

Ostapcuk, V. et al. (2018) “Activity-dependent neuroprotective protein recruits HP1 and CHD4 to control lineage-specifying genes,” Nature, 557(7707), pp. 739–743.

Padeken, J., Methot, S.P. and Gasser, S.M. (2022) “Establishment of H3K9-methylated heterochromatin and its functions in tissue differentiation and maintenance,” Nature Reviews. Molecular Cell Biology, 23(9), pp. 623–640.

Peric-Hupkes, D. et al. (2010) “Molecular maps of the reorganization of genome-nuclear lamina interactions during differentiation,” Molecular Cell, 38(4), pp. 603–613.

Phillips, J.E. and Corces, V.G. (2009) “CTCF: master weaver of the genome,” Cell, 137(7), pp. 1194–1211.

Ritchie, M.E. et al. (2015) “limma powers differential expression analyses for RNA-sequencing and microarray studies,” Nucleic Acids Research, 43(7), p. e47.

Rossi, M.J. et al. (2021) “A high-resolution protein architecture of the budding yeast genome,” Nature, 592(7853), pp. 309–314.

Rossi, M.J., Lai, W.K.M. and Pugh, B.F. (2018) “Simplified ChIP-exo assays,” Nature communications, 9(1). Available at: 10.1038/s41467-018-05265-7.

Rullens, P. et al. (2025) “MeCP2 binding and genome–lamina reorganization precede long gene activation during mouse corticogenesis,” bioRxiv. Available at: 10.1101/2025.09.01.673434.

Satterstrom, F.K. et al. (2020) “Large-Scale Exome Sequencing Study Implicates Both Developmental and Functional Changes in the Neurobiology of Autism,” Cell, 180(3), pp. 568–584.e23.

Schaefer, A. (2012) “Control of neuronal gene transcription and behavior by the epigenetic suppressor complex G9a/GLP,” in Research and Perspectives in Neurosciences. Berlin, Heidelberg: Springer Berlin Heidelberg (Research and Perspectives in Neurosciences), pp. 63–70.

Skene, P.J. and Henikoff, S. (2017) “An efficient targeted nuclease strategy for high-resolution mapping of DNA binding sites,” eLife, 6. Available at: 10.7554/eLife.21856.

Song, M. et al. (2015) “Slitrk5 Mediates BDNF-Dependent TrkB Receptor Trafficking and Signaling,” Developmental cell, 33(6), pp. 690–702.

van Steensel, B. and Belmont, A.S. (2017) “Lamina-associated domains: Links with chromosome architecture, heterochromatin, and gene repression,” Cell, 169(5), pp. 780–791.

Stessman, H.A.F. et al. (2016) “Disruption of POGZ Is Associated with Intellectual Disability and Autism Spectrum Disorders,” American journal of human genetics, 98(3), pp. 541–552.

Sun, X. et al. (2023) “POGZ suppresses 2C transcriptional program and retrotransposable elements,” Cell reports, 42(8), p. 112867.

Sun, X., Cheng, L. and Sun, Y. (2022) “Autism-associated protein POGZ controls ESCs and ESC neural induction by association with esBAF,” Molecular autism, 13(1), p. 24.

Tachibana, M. et al. (2005) “Histone methyltransferases G9a and GLP form heteromeric complexes and are both crucial for methylation of euchromatin at H3-K9,” Genes & Development, 19(7), pp. 815–826.

Xu, S. et al. (2014) “Spatial clustering for identification of ChIP-enriched regions (SICER) to map regions of histone methylation patterns in embryonic stem cells,” *Methods in molecular biology (Clifton*, N.J.), 1150, pp. 97–111.

