## supplementary figures for "POGZ safeguards neuronal gene chromatin architecture and transcription"

**Figure S1**

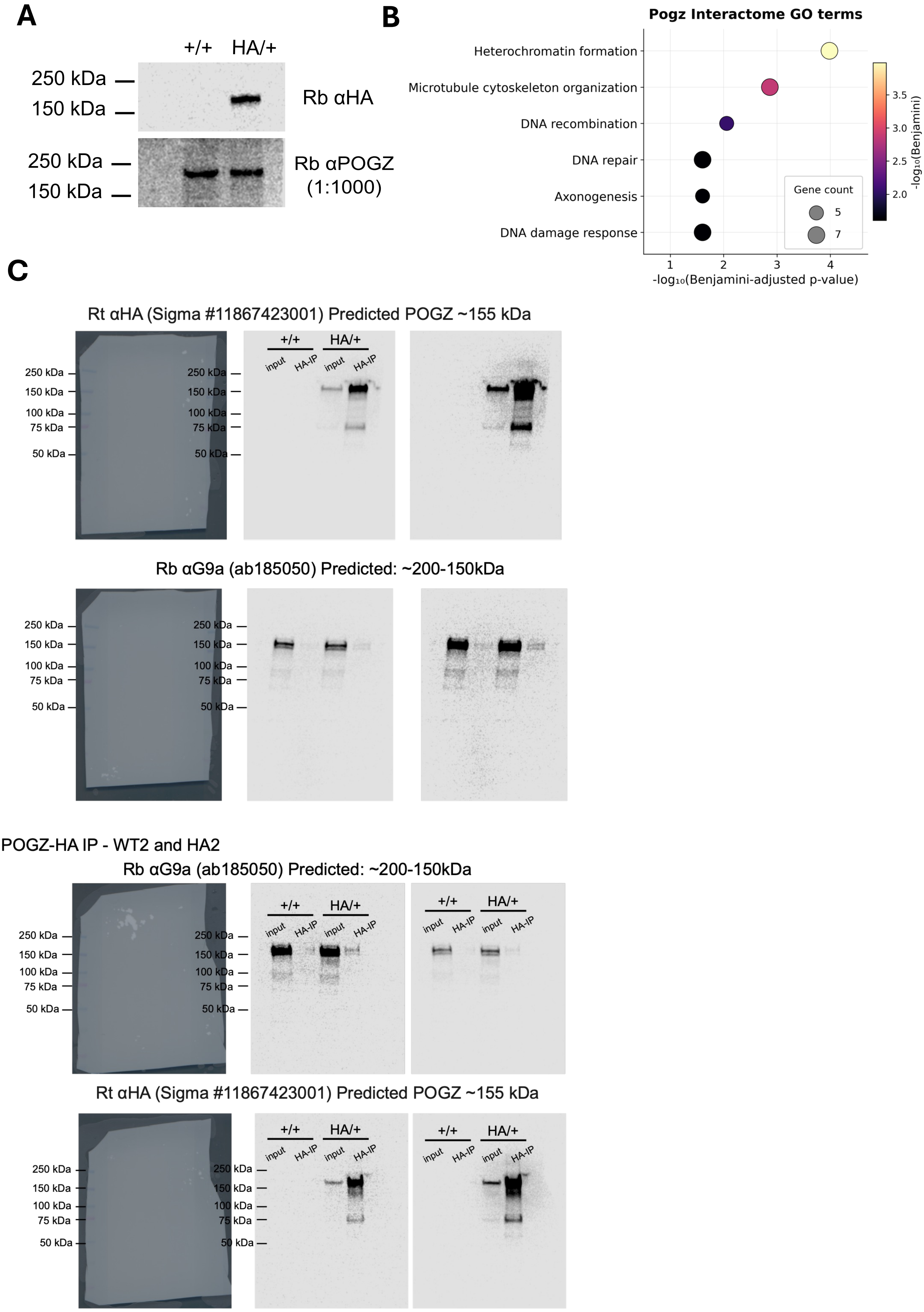
A
+/+
HA/+
250 kDa
150 kDa
Rb αHA
250 kDa
150 kDa
Rb αPOGZ
(1:1000)
B
Pogz Interactome GO terms
Heterochromatin formation
Microtubule cytoskeleton organization
DNA recombination
DNA repair
Axonogenesis
DNA damage response
-log₁₀(Benjamini-adjusted p-value)
-log₁₀(Benjamini)
Gene count
5
7
C
Rt αHA (Sigma #11867423001) Predicted POGZ ~155 kDa
input
HA-IP
250 kDa
150 kDa
100 kDa
75 kDa
50 kDa
Rb αG9a (ab185050) Predicted: ~200-150kDa
POGZ-HA IP - WT2 and HA2
Rb αG9a (ab185050) Predicted: ~200-150kDa
Rt αHA (Sigma #11867423001) Predicted POGZ ~155 kDa

**Figure S1. IP-MS and co-IP with Pogz-HA knock-in mouse**

(A) Western blot validation of the Pogz-HA knock-in mouse. Anti-HA immunoblot of E13.5 liver lysate from a *Pogz* $^{+/HA}$ heterozygous knock-in embryo and a wild-type littermate. A single HA-reactive band at the expected size for POGZ is detected in the *Pogz-HA* lane and is absent in wild-type (top). Immunoblotting with a POGZ antibody shows bands in both lanes, confirming POGZ expression in both genotypes (bottom).

(B) Gene Ontology enrichment of proteins enriched in anti-HA IP over wild-type ($log_2FC$ > 0.5). Dot size, number of proteins; color, enrichment significance.

(C) Co-immunoprecipitation of POGZ and G9a in the developing mouse cortex. HA and G9a expression in wild-type (WT-1, WT-2) and *Pogz*-HA heterozygous (HA-1, HA-2) cortices, with 5% input (input, 20 ug total protein) or after HA pull down (HA-IP). G9a blots shown are the same blots with higher and lower exposure settings.

### Figure S2

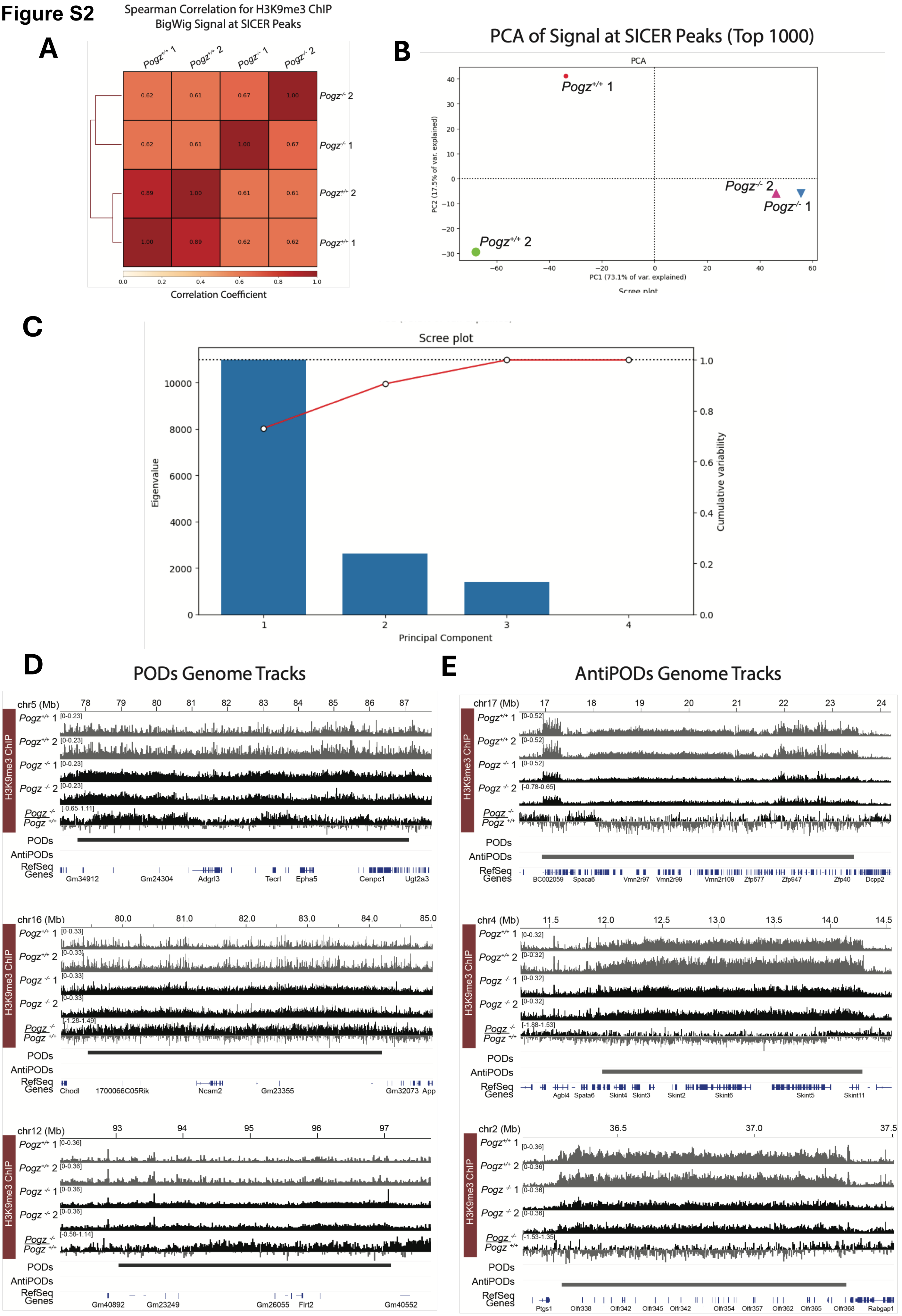
A
Spearman Correlation for H3K9me3 ChIP BigWig Signal at SICER Peaks
Correlation Coefficient
B
PCA of Signal at SICER Peaks (Top 1000)
PC1 (73.1% of var. explained)
PC2 (17.5% of var. explained)
C
Scree plot
Eigenvalue
Cumulative variability
Principal Component
D
PODs Genome Tracks
E
AntiPODs Genome Tracks
H3K9me3 ChIP
PODs
AntiPODs
RefSeq Genes

**Figure S2. Quality control of H3K9me3 ChIP-seq and genome-wide POD/anti-POD tracks.**

(A) Spearman correlation matrix of H3K9me3 ChIP-seq signal (BigWig) at SICER peaks across replicates (*Pogz*+/+ 1, 2; *Pogz* $^{-/-}$ 1, 2).

(B) Principal component analysis of H3K9me3 signal at the top 1,000 SICER peaks, colored by genotype.

(C) Variance explained per principal component (scree plot).

(D) Genome browser tracks of H3K9me3 (per replicate and *Pogz* $^{-/-}$ normalized to WT) at representative POD loci.

(E) As in (D) for representative anti-POD loci.

**Figure S3**

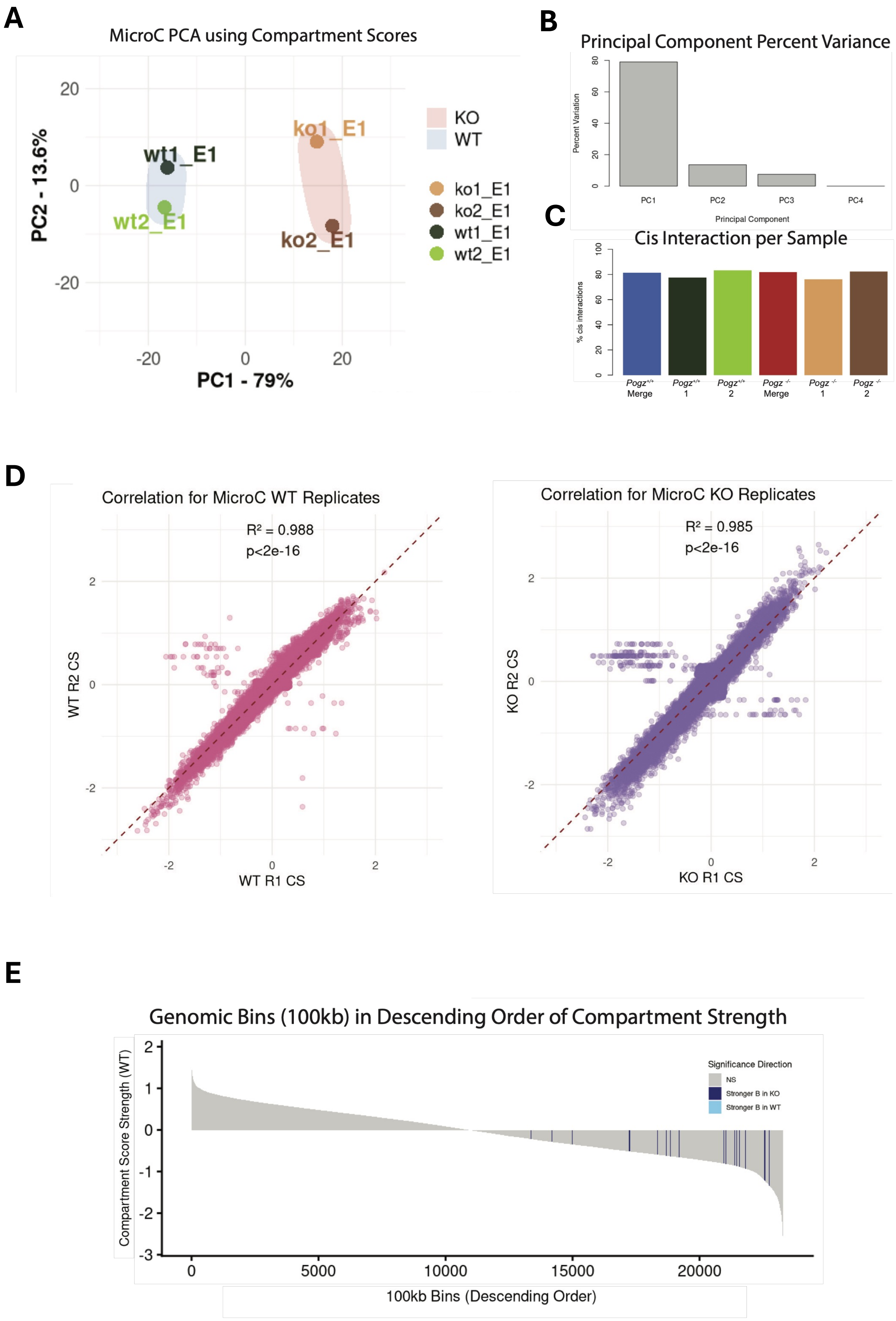
A
MicroC PCA using Compartment Scores
PC2 - 13.6%
PC1 - 79%
ko1_E1
ko2_E1
wt1_E1
wt2_E1
KO
WT
B
Principal Component Percent Variance
Percent Variation
Principal Component
PC1
PC2
PC3
PC4
C
Cis Interaction per Sample
% cis interactions
Pogz+/+ Merge
Pogz+/+ 1
Pogz+/+ 2
Pogz-/- Merge
Pogz-/- 1
Pogz-/- 2
D
Correlation for MicroC WT Replicates
R² = 0.988
p<2e-16
WT R2 CS
WT R1 CS
Correlation for MicroC KO Replicates
R² = 0.985
p<2e-16
KO R2 CS
KO R1 CS
E
Genomic Bins (100kb) in Descending Order of Compartment Strength
Compartment Score Strength (WT)
100kb Bins (Descending Order)
Significance Direction
NS
Stronger B in KO
Stronger B in WT

**Figure S3. Quality control and statistical validation of Micro-C compartment analysis.**

**(A)** PCA of Micro-C compartment scores; replicates (ko1, ko2, wt1, wt2) cluster by genotype.

**(B)** Percent variance explained per principal component.

**(C)** Percentage of cis interactions per sample.

**(D)** Correlation of compartment scores between Micro-C replicates: wild-type ($R^2 = 0.988$) and *Pogz*$^{-/-}$ ($R^2 = 0.985$); $p < 2 \times 10^{-16}$.

**(E)** Compartment strength scores across 100-kb genomic bins ranked in descending order, with significantly changed bins annotated by direction (stronger B in KO; stronger B in WT).

### Figure S4

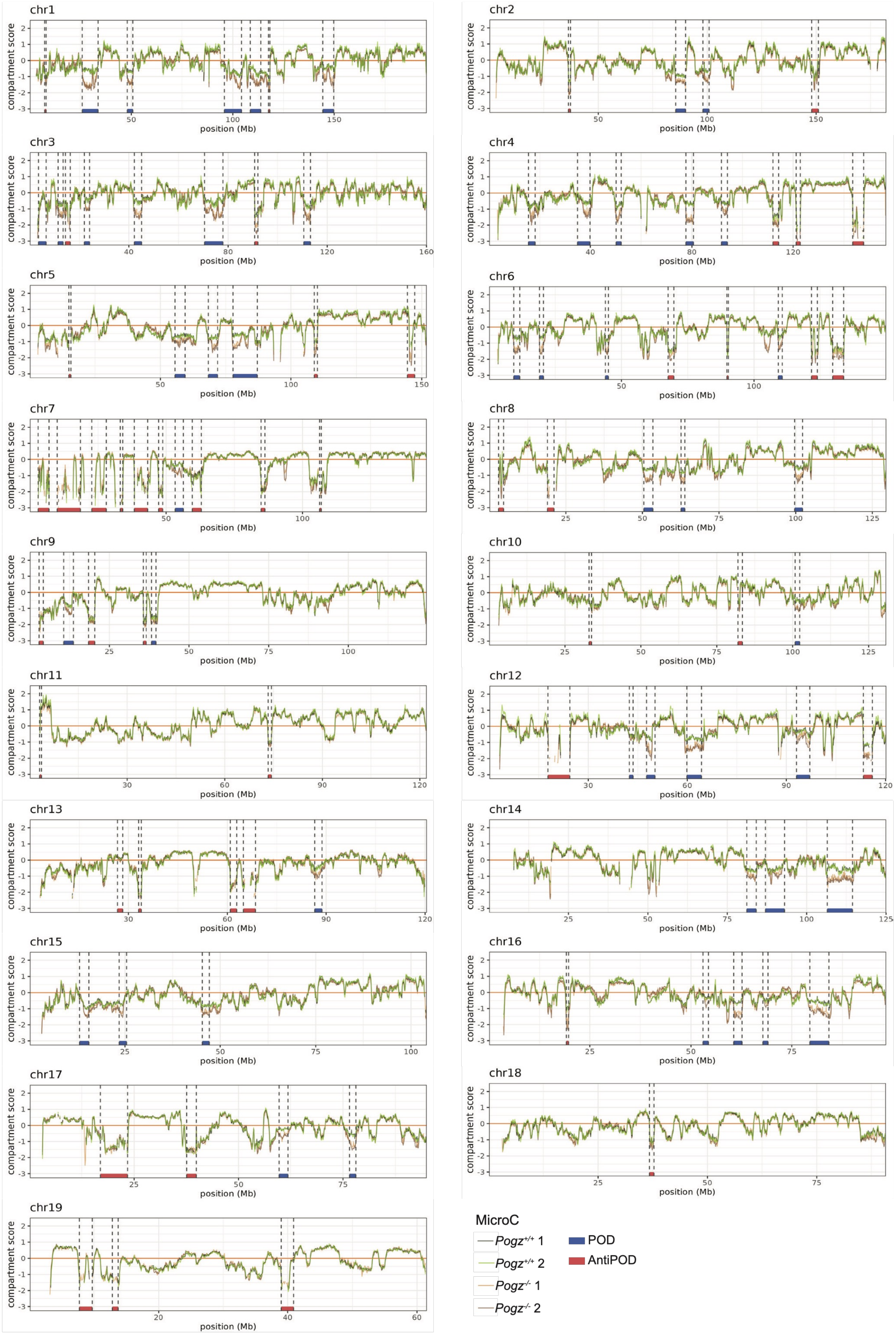
chr1
chr2
chr3
chr4
chr5
chr6
chr7
chr8
chr9
chr10
chr11
chr12
chr13
chr14
chr15
chr16
chr17
chr18
chr19
compartment score
position (Mb)
MicroC
Pogz+/+ 1
Pogz+/+ 2
Pogz-/- 1
Pogz-/- 2
POD
AntiPOD

**Figure S4. Genome-wide Micro-C compartment strength score tracks.**

Per-chromosome Micro-C compartment strength score tracks for WT(rep1 & rep 2) and *Pogz* $^{-/-}$ (rep1 & rep2). POD (blue) and anti-POD (red) intervals annotated below each track.

**Figure S5**

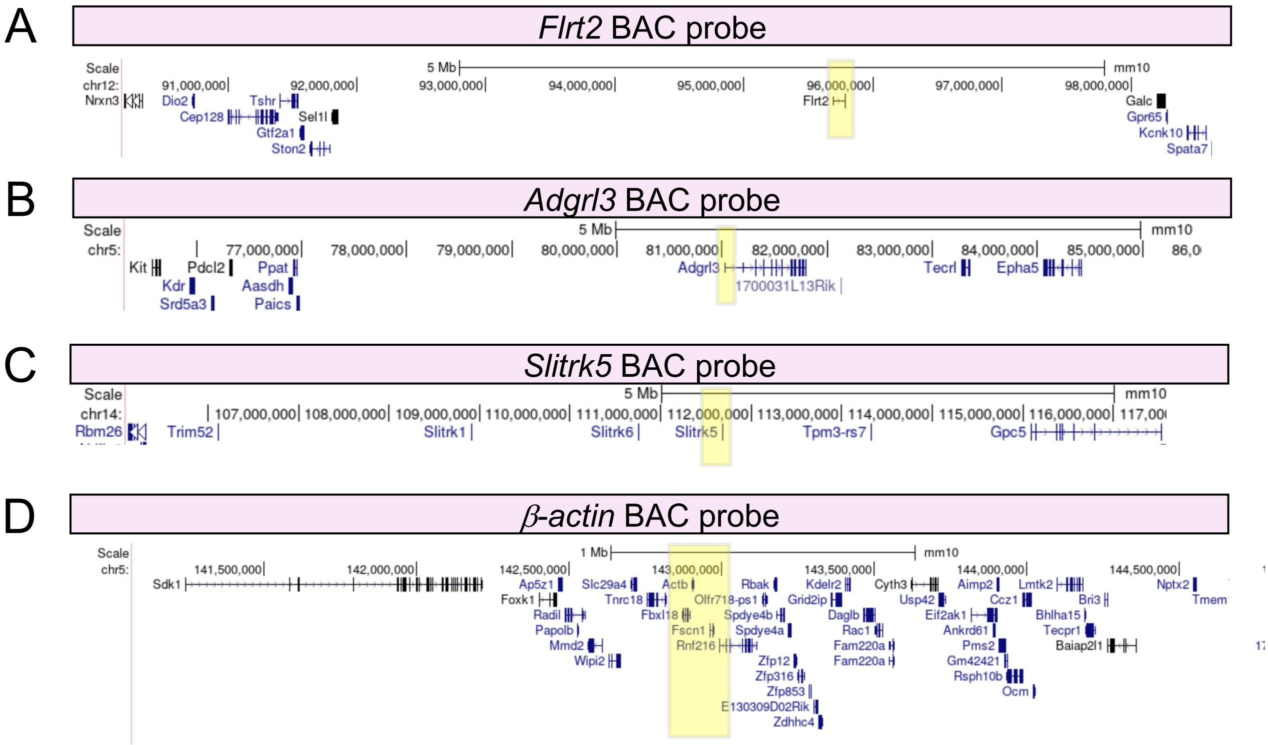

A
Flrt2 BAC probe
Scale
5 Mb
mm10
chr12:
91,000,000
92,000,000
93,000,000
94,000,000
95,000,000
96,000,000
97,000,000
98,000,000
Nrxn3
Dio2
Tshr
Cep128
Sel1l
Gtf2a1
Ston2
Flrt2
Galc
Gpr65
Kcnk10
Spata7
B
Adgrl3 BAC probe
Scale
5 Mb
mm10
chr5:
77,000,000
78,000,000
79,000,000
80,000,000
81,000,000
82,000,000
83,000,000
84,000,000
85,000,000
Kit
Pdcl2
Ppat
Kdr
Aasdh
Srd5a3
Paics
Adgrl3
1700031L13Rik
Tecrl
Epha5
C
Slitrk5 BAC probe
Scale
5 Mb
mm10
chr14:
107,000,000
108,000,000
109,000,000
110,000,000
111,000,000
112,000,000
113,000,000
114,000,000
115,000,000
116,000,000
Rbm26
Trim52
Slitrk1
Slitrk6
Slitrk5
Tpm3-rs7
Gpc5
D
β-actin BAC probe
Scale
1 Mb
mm10
chr5:
141,500,000
142,000,000
142,500,000
143,000,000
143,500,000
144,000,000
144,500,000
Sdk1
Ap5z1
Foxk1
Radil
Papolb
Mmd2
Wipi2
Slc29a4
Tnrc18
Fbxl18
Actb
Olfr718-ps1
Fscn1
Rnf216
Rbak
Spdye4b
Spdye4a
Zfp12
Zfp316
Zfp853
E130309D02Rik
Zdhhc4
Kdelr2
Grid2ip
Daglb
Rac1
Fam220a
Cyth3
Usp42
Eif2ak1
Ankrd61
Pms2
Gm42421
Rsph10b
Ocm
Aimp2
Ccz1
Lmtk2
Bri3
Bhlha15
Tecpr1
Baiap2l1
Nptx2

**Figure S5. BAC probe locations for DNA FISH.**

Genomic locations (mm10) of BAC probes used for DNA FISH, highlighted in yellow, with RefSeq gene annotations: **(A)** *Flrt2* (chr12), **(B)** *Adgrl3* (chr5), **(C)** *Slitrk5* (chr14), and **(D)** *β-actin* / *Actb* control (chr5).

Figure S6

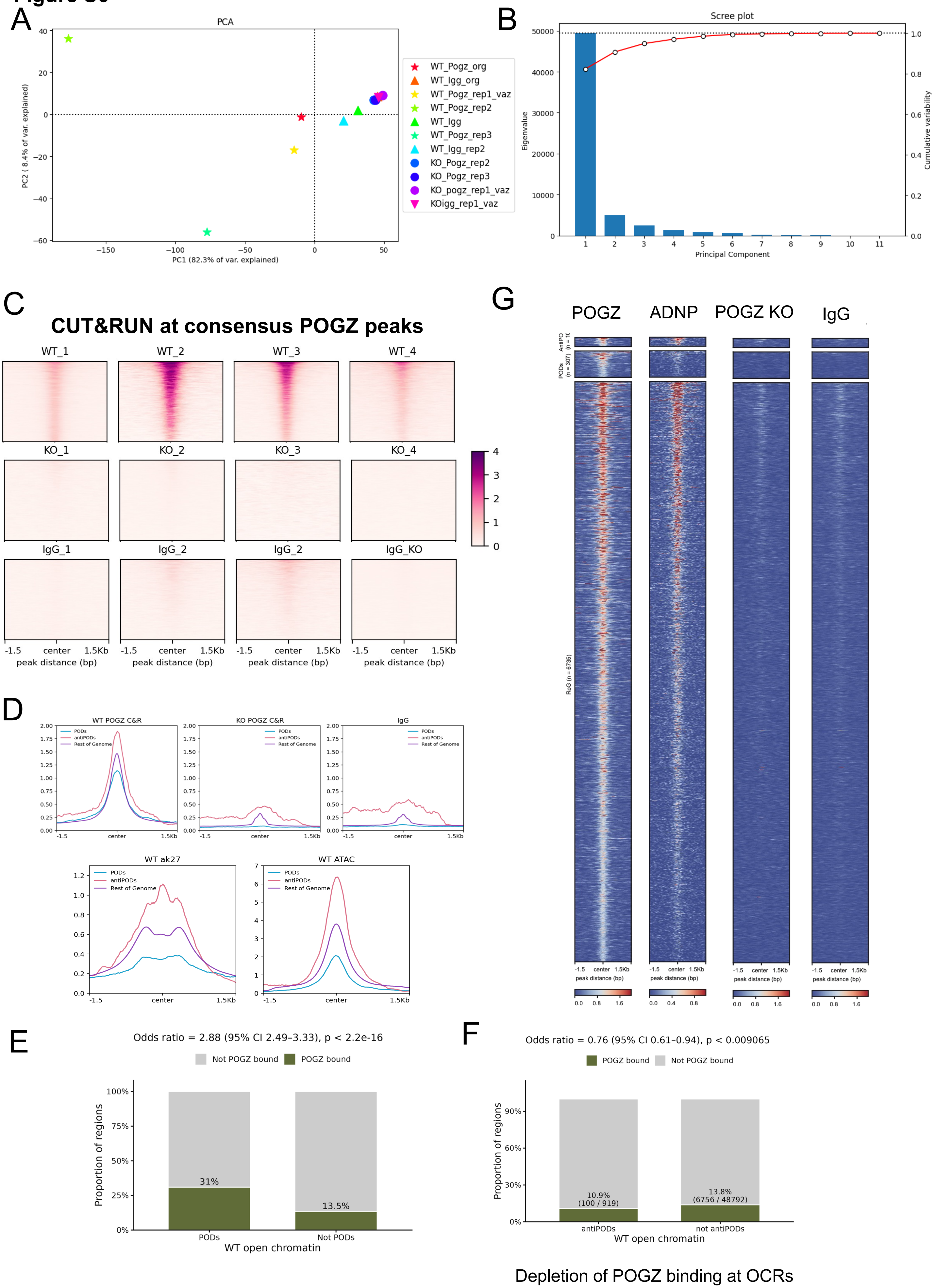


Enrichment of POGZ binding at OCRs In PODs

Depletion of POGZ binding at OCRs In anti-PODs

**Figure S6. Quality control of POGZ CUT&RUN and enrichment of POGZ binding at PODs and differentially transcribed genes.**

**(A)** PCA of POGZ CUT&RUN replicates.

**(B)** Variance explained per principal component.

**(C)** Heatmaps of POGZ CUT&RUN signal at consensus POGZ peaks (±1.5 kb) across wild-type replicates, *Pogz* $^{-/-}$ replicates, and IgG controls.

**(D)** Aggregate POGZ CUT&RUN signal profiles across the same samples.

**(E)** Proportion of wild-type open chromatin regions (OCRs) bound by POGZ within PODs versus outside PODs (31% vs 13.5%; odds ratio = 2.88, 95% CI 2.49–3.33, $p < 2.2 \times 10^{-16}$, Fisher's exact test).

**(F)** As in (E) for anti-PODs (10.9% [100/919] vs 13.8% [6,756/48,792]; odds ratio = 0.76, 95% CI 0.61–0.94, $p < 0.01$).

**(G)** Profile plots and heatmaps of CUT&RUN signal at merged POGZ peaks, split into PODs, anti-PODs, and rest of genome. Columns: POGZ (WT), ADNP, POGZ (KO control), IgG. ADNP tracks POGZ genome-wide but drops out within PODs and anti-PODs, indicating POGZ binds these domains independently of ADNP/ChAHP. Signal plotted ±1 kb from peak center.

### Figure S7

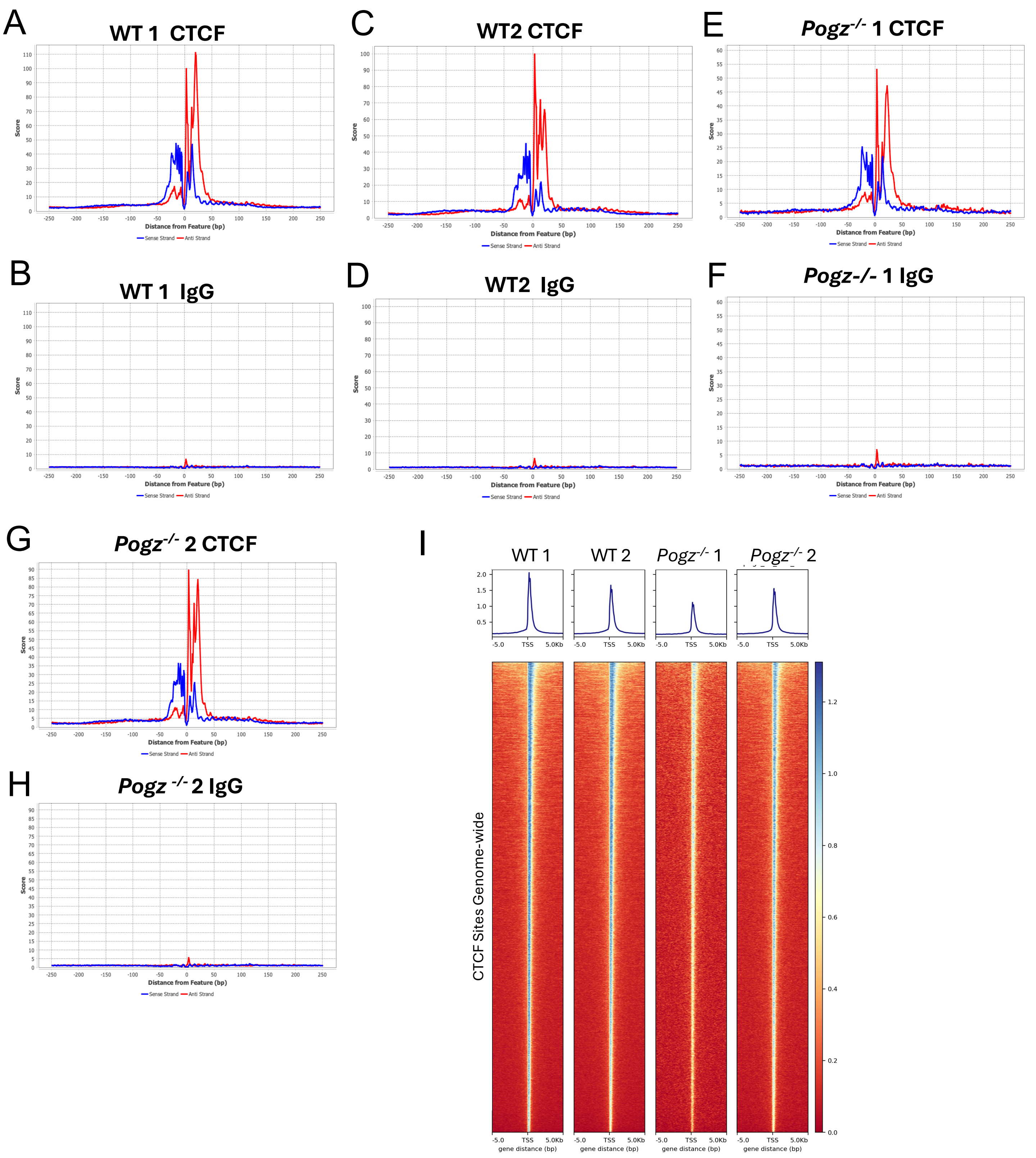

A
WT 1 CTCF
B
WT 1 IgG
C
WT2 CTCF
D
WT2 IgG
E
Pogz-/- 1 CTCF
F
Pogz-/- 1 IgG
G
Pogz-/- 2 CTCF
H
Pogz -/- 2 IgG
Score
Distance from Feature (bp)
Sense Strand
Anti Strand
I
WT 1
WT 2
Pogz-/- 1
Pogz-/- 2
TSS
5.0Kb
-5.0
gene distance (bp)
CTCF Sites Genome-wide

**Figure S7. Quality control of CTCF ChIP-exo.**

**(A-H)** CTCF ChIP-exo signal pileup at CTCF motifs for wild-type cortex and in *Pogz* $^{-/-}$ replicates and IgG controls demonstrating near base-pair resolution.

**(E)** Aggregate CTCF ChIP-exo signal for WT and *Pogz* $^{-/-}$ replicates normalized (IgG normalized) centered on CTCF sites genome-wide.

**Figure S8**

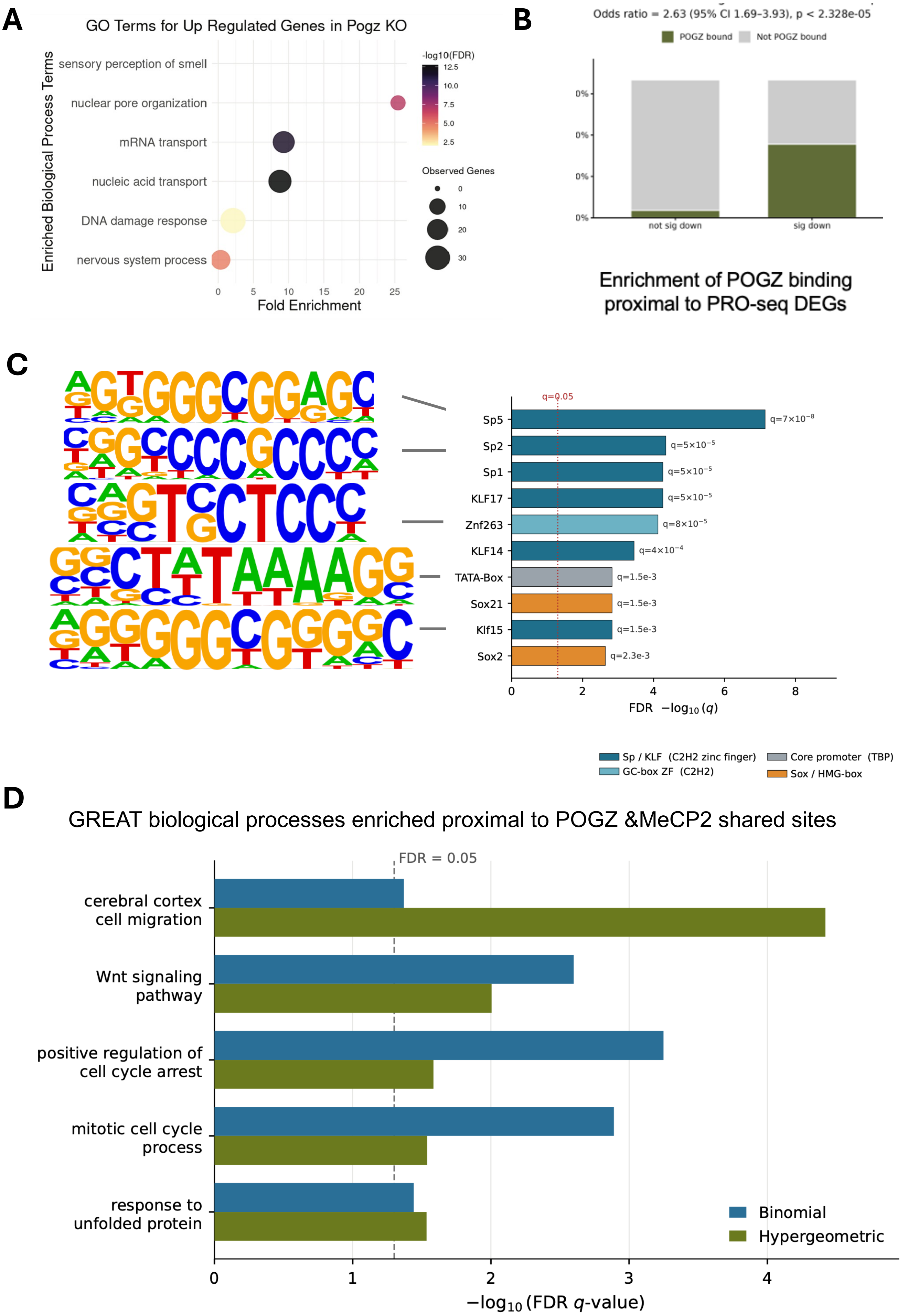
A
GO Terms for Up Regulated Genes in Pogz KO
Enriched Biological Process Terms
sensory perception of smell
nuclear pore organization
mRNA transport
nucleic acid transport
DNA damage response
nervous system process
Fold Enrichment
-log10(FDR)
Observed Genes
B
Odds ratio = 2.63 (95% CI 1.69–3.93), p < 2.328e-05
POGZ bound
Not POGZ bound
not sig down
sig down
Enrichment of POGZ binding proximal to PRO-seq DEGs
C
q=0.05
Sp5 q=7×10⁻⁸
Sp2 q=5×10⁻⁵
Sp1 q=5×10⁻⁵
KLF17 q=5×10⁻⁵
Znf263 q=8×10⁻⁵
KLF14 q=4×10⁻⁴
TATA-Box q=1.5e-3
Sox21 q=1.5e-3
Klf15 q=1.5e-3
Sox2 q=2.3e-3
FDR −log10 (q)
Sp / KLF (C2H2 zinc finger)
GC-box ZF (C2H2)
Core promoter (TBP)
Sox / HMG-box
D
GREAT biological processes enriched proximal to POGZ &MeCP2 shared sites
FDR = 0.05
cerebral cortex cell migration
Wnt signaling pathway
positive regulation of cell cycle arrest
mitotic cell cycle process
response to unfolded protein
Binomial
Hypergeometric
−log10 (FDR q-value)

**Figure S8. GO enrichment of up-regulated genes and POGZ binding proximal to down-regulated genes in *Pogz*$^{-/-}$ cortex.**

**(A)** GO Biological Process terms enriched among genes with significantly increased nascent transcription (PRO-seq) in *Pogz*$^{-/-}$ versus wild-type E13.5 cortex. Dot size, number of observed genes; color, $-\log_{10}$(FDR); x-axis, fold enrichment.

**(B)** Proportion of genes with POGZ CUT&RUN binding (green) versus no binding (gray) among genes that are not significantly down-regulated ("not sig down") and significantly down-regulated ("sig down") in *Pogz*$^{-/-}$ cortex. Down-regulated genes are enriched for proximal POGZ binding (odds ratio = 2.63, 95% CI 1.69–3.93, $p < 2.33 \times 10^{-5}$, Fisher's exact test).

**(C)** HOMER known-motif enrichment at POGZ CUT&RUN peaks in promoters of genes down-regulated in POGZ KO. Top 10 motifs ranked by Benjamini–Hochberg FDR ($-\log_{10}$ q; dotted line, q = 0.05), colored by TF family. Logos at left show representative motifs.

**(D)** GREAT enrichment (mm10, single nearest gene). Bars show $-\log_{10}$(FDR q-value) for the binomial (blue) and hypergeometric (green) tests; dashed line marks FDR = 0.05. All terms are significant in both tests, ordered by hypergeometric significance.
